# Increased juvenile survival may not be universally linked to longevity: ecological, social and life-history drivers of age-specific mortality in birds

**DOI:** 10.1101/2019.12.18.880682

**Authors:** Emeline Mourocq, Szymon M. Drobniak, Michael Griesser

## Abstract

A classical prediction of the traditional evolutionary theories of ageing (tETA) is that longevity should be positively correlated with survival early on in life. However, large and unexplained variation exists in juvenile survival-longevity combinations. Here, we provide the first comparative study investigating the life-history, ecological and social correlates of juvenile survival, longevity and their combinations in 204 bird species. Overall, both measurements were positively correlated, but multiple survivals’ combinations evolved, some in accordance with tETA (“positive JS-L combinations”) while others contrasting it (“JS-L mismatches”). Positive JS-L combinations covaried with the pace of life proxies, whereas mismatching combinations covaried with the growing season length, where long growing seasons promoted juvenile survival, while short growing seasons promoted longevity. Interestingly, sociality explained only positive combinations, while life-history and ecological traits explained both positive and mismatching combinations. Overall, these findings challenge a main prediction of the tETA, and identify key evolutionary forces driving the coevolution between juvenile survival and longevity.

---

Traditional theories of aging (tETA: “mutation accumulation”^1^, “antagonistic pleiotropy”^2^, “antagonistic pleiotropy”^3^) propose that extrinsic mortality is the main driver of longevity^4,5^. They predict that higher extrinsic mortality early on in life leads to relatively few individuals reaching old age, and the fitness value of prolonged lifespan is therefore small in such cases. Thus, selection to extend longevity is only strong in populations with high survival early on in life (juvenile survival henceforth). Accordingly, these theories predict that longevity should be positively correlated with juvenile survival^2,4,6–8^.

Although this classical prediction of tETA underlies many life-history studies, and is commonly cited as being largely corroborated by existing data^9,10^, support has been mixed and alternative theories exist^5,11–13^. Moreover, recent theoretical and empirical studies do challenge this prediction^9,14–16^, and state that juvenile survival (extrinsic mortality early on in life) is not a random process but does depend on age, individual condition, or population density. Accordingly, species can deviate from the expected relationship between juvenile survival and longevity, by having a low juvenile survival but being long-lived, or by having a high juvenile survival but being short-lived^5,9,13–17^. However, it remains unclear whether these deviations represent evolved strategies modulated by specific life-history, ecological and/or social factors, or whether they are pieces of a continuum of randomly varying combinations.

Longevity is a pivotal factor shaping life-histories^18,19^, but survival can vary among the stages of life and differently influence the evolution of life-history traits^20–22^. Theoretical^12,23–26^ and empirical work on birds^27^, fishes^22^ and mammals^28^ have highlighted the importance of considering age-specific survival to understand the evolution of life-history traits. Specifically, these studies showed that agespecific survival patterns that deviate from the classical prediction of tETA (e.g., low chance of survival early in life but a high longevity), are linked to unusual combinations of life-history traits that are characteristic to both slow- and fast-living animals^19^. For instance, turtles and crocodiles suffer from high juvenile mortality, and accordingly females lay many eggs in each reproductive event (like fast-living animals) despite that they are exceptionally long-lived (like slow-living animals)^29^. Thus, considering the factors affecting age-specific survivals and their combination is critical to understand life history evolution in general.

Longevity varies considerably across species. In vertebrates it ranges from a few months to over 100 years^30^. Comparative work did show that adaptations that reduce extrinsic mortality, including protective shells^6^ or the ability to fly^31^, are linked with increased longevity. Moreover, long-lived species tend to be active during the period of day with the lowest predation risk^31^, have a low number of co-occurring predators of adults^32^ or life-history traits characteristic of a slow pace of life (e.g., produce few offspring, which develop slowly and mature relatively late in life)^12,18^. Additionally, larger mammals and birds live longer than smaller ones^32–34^.

In many taxa, juveniles usually have lower and more variable survival than adults^35–38^. The few studies investigating juvenile survival showed that the small body size of juveniles may explain their low survival in lineages with slow growth (mammals, reptiles) and indeterminate growth (fish)^39,40^. In lineages with rapid body growth (birds), low juvenile survival can reflect age-dependent social dominance^35^ or lacking skills^41^. Besides, juvenile survival tends to be high in birds with long nestling periods^42^, low reproductive allocation^43,44^, prolonged post-fledging care^45^, or prolonged association with the parents beyond independence (i.e., family-living species, see^46^)^47,48^ (Table 1). Although a number of studies have investigated inter-specific variation in longevity^31,32,34,49,50^, it is unknown which factors influence survival early on in life and how this relates to longevity^51^. Importantly, comparative studies are lacking.

**Table 1.**
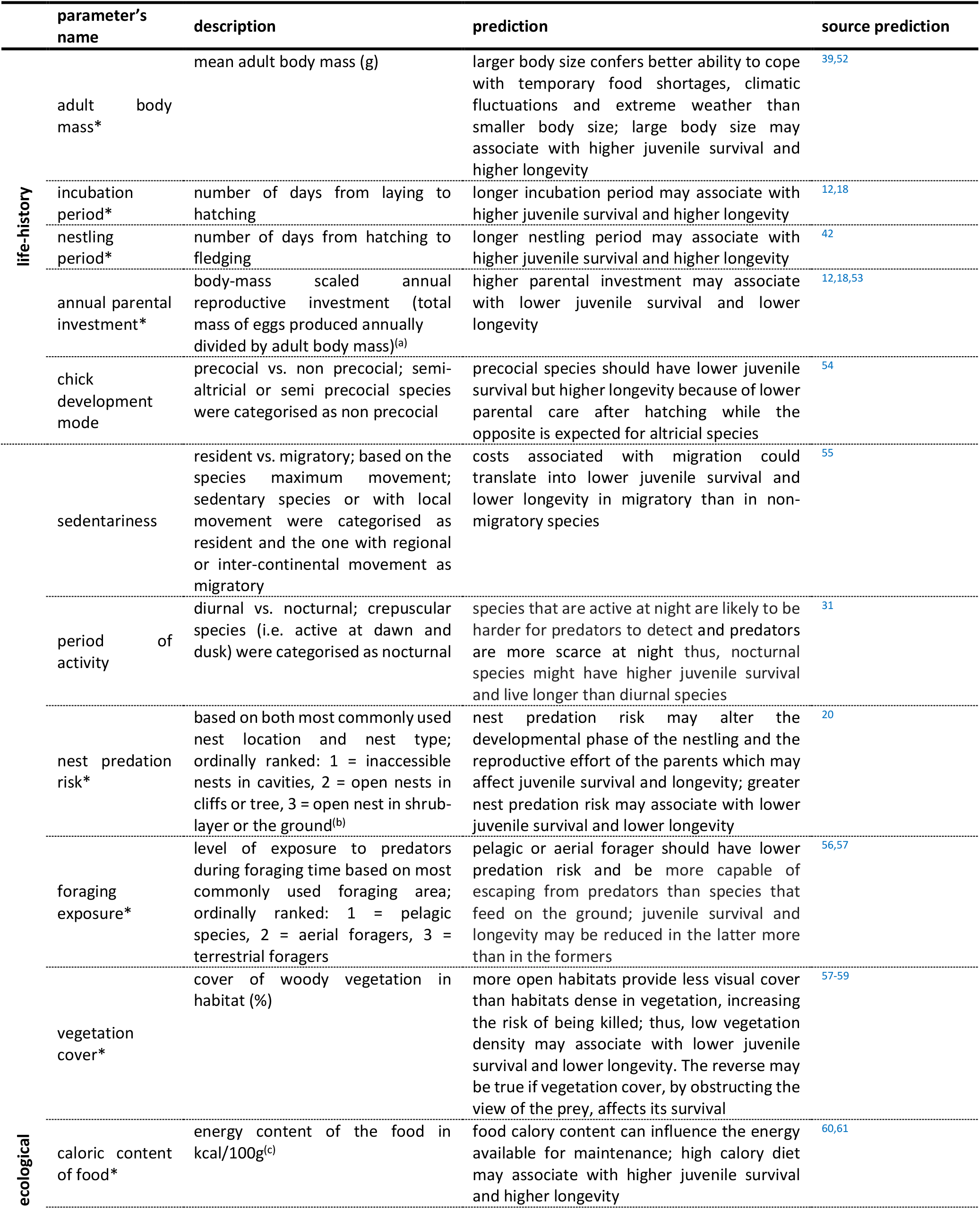
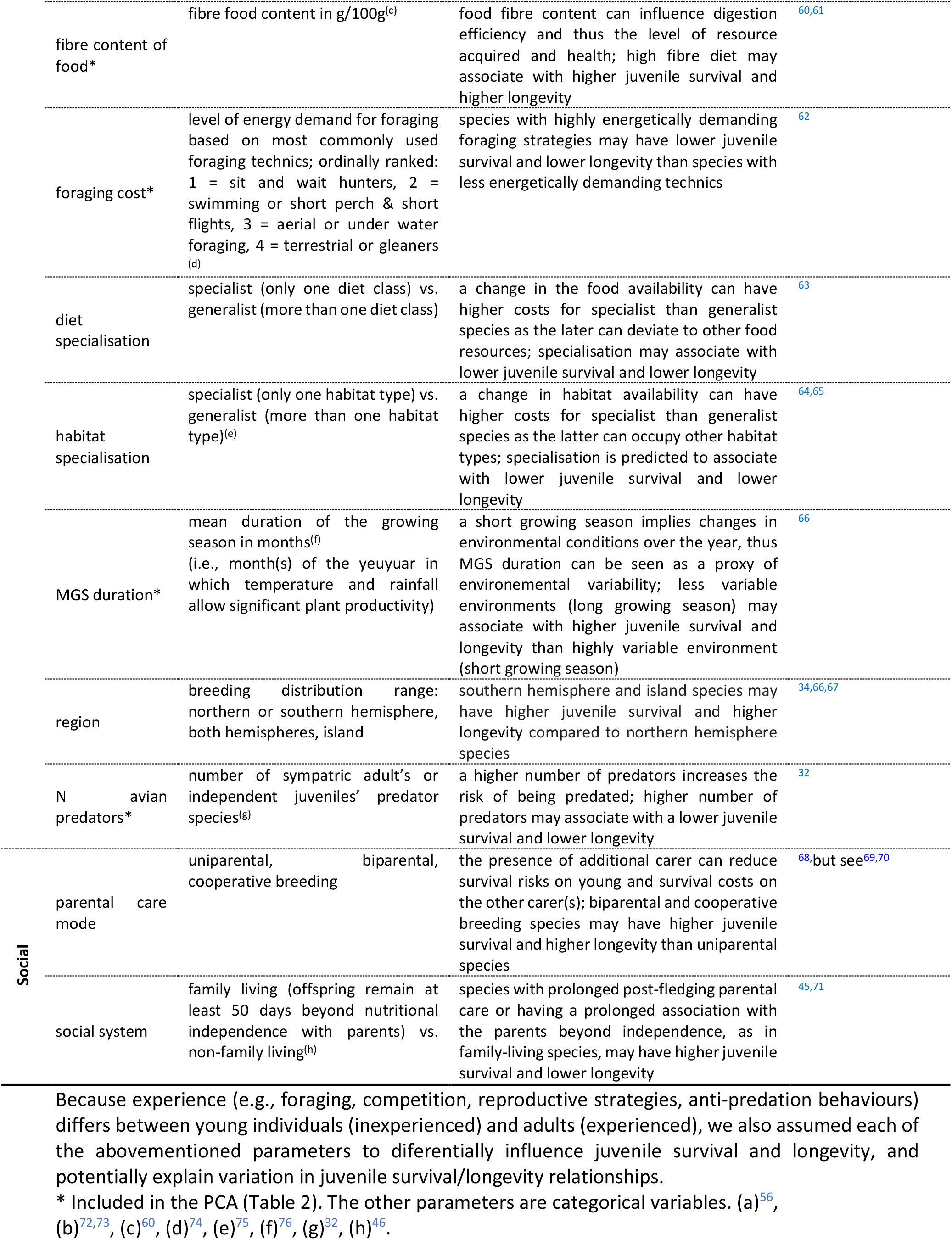
Description and prediction of the parameters investigated in this study.

Here, we use phylogenetic comparative analyses to understand interspecific variation in juvenile survival (measured as post-fledglings to first-year survival rate) and maximum longevity, as well as their relationship, in 204 bird species. Firstly, we compare the association of (i) juvenile survival and (ii) maximum longevity with life-history, ecological and social parameters. Secondly, we investigate how juvenile survival and maximum longevity relate to each other, and assess which lifehistory, ecology and social traits better explain (i) positive associations between juvenile survival and longevity (i.e., as expected by the classical prediction of tETA: low-low and high-high combinations, referred to as “positive JS-L combinations” henceforth), and (ii) mismatches between juvenile survival and longevity (i.e., deviation from the classical prediction of tETA: low-high and high-low combinations, referred to as “JS-L mismatches” henceforth).

## Materials and Methods

### Survival data

We collected data on juvenile survival and maximum longevity for 293 bird species covering 20 taxonomic orders and 74 families (Fig. S1 in Supporting Information), using existing datasets^32^, the Handbook of the Birds of the World^77^, the Birds of North America^78^, the Handbook of Australian, New Zealand and Antarctic Birds^79^, the Handbook of Southern Africa^80^, the Australian Birds and Bats Banding Scheme database^81^ and Animal Ageing and Longevity database^82^ (available at http://genomics.senescence.info/species/).

Juvenile survival was assessed as the proportion of fledglings that survive their first year of life, where many juveniles die due to extrinsic mortality^83^. For species where multiple values of juvenile survival were available we used their mean. Maximum longevity (maximum observed lifespan) was mostly assessed with mark-recapture of ringed wild birds, but for 19 species longevity was of unknown origin (captivity or wild). Earlier studies showed that longevity records in captivity and the wild are highly correlated^32,34^ and thus, we also included longevity data of unknown origin. Longevity estimates are influenced by the sampling effort because the larger the sample the higher is the chance to sample a long-lived indvidual^32^. Therefore, to adjust for any bias associated with maximum longevity estimates we included the independent number of Web of Science records per species (research effort) as a covariate in our analyses (available at http://apps.webofknowledge.com).

### Life-history, ecology and social parameters

We used a published dataset^84^ that was complemented with data from the sources listed above, and compiled data on life-history, ecological and social parameters that may influence juvenile survival and longevity (Table 1). We could find data for the 20 parameters listed in Table 1 for 204 of the 293 species (Fig. S2). Thus, 293 species were considered in descriptive analyses, while a subset 204 species entered detailed phylogenetic mixed models.

### Statistical analyses

#### General procedures

All statistical analyses were performed in R version 3.2.2^85^. We used phylogenetic controlled mixed models in ASReml-R 3^86^ to control for the phylogenetic dependency among species (VSN International, Hempstead, U.K.^87^). We included phylogeny as a random effect in the model in the form of a correlation matrix of distances from the root of the tree to the most recent common ancestor between two species. We tested the phylogenetic effect with a likelihood ratio test where 2 times the difference in log-likelihood between the model with and without the phylogeny is tested against a χ^2^ distribution with one degree of freedom^88^. To account for phylogenetic uncertainty, all ASReml-R models were run with 300 different phylogenetic trees obtained from www.birdtree.org^89^. We averaged the estimates from the 300 models and present the averaged estimates and the Fs300 (proportion of trees for which the p-value associated with an estimate was <0.05). Individual p-values were obtained through a conditional Wald F-test. All continuous variables were standardised by centring (around the mean) and scaling (by the standard deviation) them, to allow direct comparison of the model estimates^90^, but we present raw data in the figures. We checked for the assumptions of normally distributed and homogeneous residuals by visually inspecting histograms and qq-plots of the residuals as well as residuals plotted against fitted values.

To reduce the multidimensionality of our predictor variables and to reduce their collinearity^91^, we performed a principal component analysis (PCA) with varimax rotation including all 12 continuous predictors, and extracted 7 PC’s given in Table 2. Prior to the PCA, the distribution of these predictors was checked graphically and, if necessary, transformed to obtain a more symmetrical distribution, and then standardised (see above).

**Table 2.** **Results of the Principal Component Analysis (PCA) with** varimax rotation on the 12 continuous predictors.

|  |  |  |  | life-history<br>pace | exposure<br>to<br>predators | food<br>fibre | nest<br>predation<br>risk | N avian<br>predators | foraging<br>cost | MGS<br>duration |  |
| --- | --- | --- | --- | --- | --- | --- | --- | --- | --- | --- | --- |
| category | transformation | variable | n° component | 1 | 7 | 2 | 6 | 3 | 5 | 4 | h2 |
| life-history | ln | adult body mass |  | **0.90** | -0.25 | -0.08 | 0.13 | -0.01 | -0.17 | -0.12 | 0.95 |
|  | none | annual parental investment |  | **-0.89** | 0.14 | -0.17 | 0.13 | 0.14 | -0.05 | -0.02 | 0.87 |
|  | sqrt | incubation period |  | **0.76** | -0.35 | 0.23 | -0.03 | -0.10 | -0.28 | -0.04 | 0.84 |
|  | none | nestling period |  | 0.51 | 0.04 | 0.44 | -0.58 | -0.13 | 0.01 | 0.20 | 0.85 |
| ecological | none | nest predation risk |  | 0.05 | -0.10 | -0.12 | **0.94** | -0.12 | 0.09 | -0.07 | 0.93 |
|  | none | foraging cost |  | -0.17 | 0.02 | -0.11 | 0.08 | -0.07 | **0.96** | -0.05 | 0.98 |
|  | none | calorie content of food |  | 0.15 | 0.49 | 0.62 | 0.02 | 0.44 | 0.01 | -0.01 | 0.83 |
|  | ln | fibre content of food |  | -0.11 | 0.16 | **-0.89** | 0.22 | -0.01 | 0.15 | -0.10 | 0.92 |
|  | none | foraging exposure |  | -0.30 | **0.86** | -0.04 | -0.01 | 0.04 | -0.02 | 0.00 | 0.83 |
|  | sqrt | vegetation cover |  | -0.21 | **0.85** | -0.01 | -0.13 | -0.08 | 0.05 | 0.11 | 0.80 |
|  | sqrt | N avian predators |  | -0.18 | -0.03 | 0.08 | -0.07 | **0.95** | -0.05 | 0.00 | 0.95 |
|  | none | MGS duration |  | -0.06 | 0.08 | 0.08 | -0.11 | -0.01 | -0.05 | **0.98** | 0.99 |
| SS loadings |  |  |  | 2.68 | 1.94 | 1.5 | 1.34 | 1.15 | 1.07 | 1.05 |  |
| cumulative variance explained (%) |  |  |  | 22 | 39 | 51 | 62 | 72 | 81 | 89 |  |

Full mixed models included the 7 PC’s (Table 2), the 8 categorical variables described in Table 1, and as covariates research effort (log transformed) and body mass (log transformed) to control for allometry^32,49^. Since the life-history pace PC was loaded by adult body mass (Table 2) and therefore partially controlling for allometry, we only included the residuals from a linear model between the natural logarithm of adult body mass and the life-history pace component as body-mass covariate. This way the presence in the model of both the life-history pace PC and the residual body mass allows to fully control for allometry.

The importance of first-year survival for fitness benefits is likely to depend on the age at first reproduction (AFR) (63.8% of the species had an AFR ≤ 1 year old, 17%]1; 2], 9.6%]2; 3] and 9.6% > 3 years old, Fig. S3). Therefore, we re-ran the PCA and all the following analyses on a subset of species for which AFR was available (N=188, Fig. S4). PCA output remained the same, and AFR loaded positively on the life-history pace PC (Table S1). The linear mixed-effects models gave qualitatively similar output (Tables S2, S3, S4) suggesting that in our set of species it is unlikely that AFR affected our analyses, and thus we present in the manuscript the analyses including all species (N=204).

#### Correlates of juvenile survival and longevity

We ran two phylogenetically controlled linear mixed-effects models including the same life-history, ecological and social predictors to assess the factors correlating with juvenile survival and with longevity. We fitted in both cases the full models (i.e., no model selection applied) to obtain comparable estimates of the same set of predictors in both models. To compare the influence of each predictor on both response variables, juvenile survival and longevity were standardised^90^.

#### Combinations of juvenile survival and longevity

The second set of analyses assessed factors that were associated with combinations of juvenile survival and longevity that (i) concurred with (positive JS-L combinations) or (ii) deviated from (JS-L mismatches) the positive correlation between juvenile survival and longevity, predicted by tETA. We captured the natural patterns of association between juvenile survival rate and maximum longevity using a PCA approach on the two log-transformed and standardised survival variables. The PCA resulted in two principal components (PCs, Table S5). Due to the properties of a PC data rotation, PC1 was loaded positively by both survival estimates (Table S5). Thus, it describes a tied link between juvenile survival and longevity, capturing patterns that concur with the classical prediction of tETA (cases positioned on PC1 represent the most typical cases of positive JS-L combinations). PC2 was loaded positively by juvenile survival rate and negatively by maximum longevity (Table S5). Being perpendicular to PC1, it captures how much a species deviates from the overall expected association, and thus, how much it deviates from the classical prediction of tETA (JS-L mismatches).

We ran two separate phylogenetically controlled linear mixed-effects models to assess the factor associated with absolute values of (i) PC1 (positive JS-L combinations) and (ii) PC2 (JS-L mismatches). We included the same set of predictors and covariates as in the full models of juvenile survival and longevity analyses, and included the sign (positive or negative) of the corresponding PC as a factor and in interaction with each predictor. The latter allowed us to assess the correlates of each possible combination of juvenile survival and longevity, i.e., to investigate how species attributes associated with (i) high juvenile survival-high longevity vs. low juvenile survival-low longevity combinations (positive JS-L combinations, analysis of PC1), and (ii) deviation towards higher juvenile survival-lower longevity vs. lower juvenile survival-higher longevity (JS-L mismatches, analysis of PC2). For both models, we used a backward model selection process. We successively removed terms with p > 0.10, starting with the highest-order interactions and following with the simple effects. We compared models including and excluding the focal predictor using *model.sel* function from the MuMIn package^92^. The decision to exclude the predictor was based on the AICc criterion using a ΔAICc (i.e., AICc_included_ –AICc_excluded_) > 2 as threshold^93^. Results of the full models are provided in Table S6 and S7.

## Results

### Correlates of juvenile survival and longevity

Juvenile survival rate ranged from 0.08 to 0.95 (0.39 ± 0.16; mean ± SD) and maximum longevity ranged from 5 to 51 (17.7 ± 9.0) years. Juvenile survival and longevity both correlated with the life-history pace PC, where species with a slow life-history pace (large body size, low annual reproductive investment, long incubation period; Table 1) had significantly higher juvenile survival and greater longevity compared to species with a fast life-history pace (small body size, high annual reproductive investment, short incubation period; Tables 1 and 3). Moreover, juvenile survival was higher in species with a high nest predation risk (open nest close to the ground or on the ground; Tables 1 and 3), while longevity was greater in species with a low exposure of adults to predators (pelagic forager, living in open habitat; Tables 1 and 3). The phylogenetic effect was only significant for longevity (Table 3).

**Table 3.**
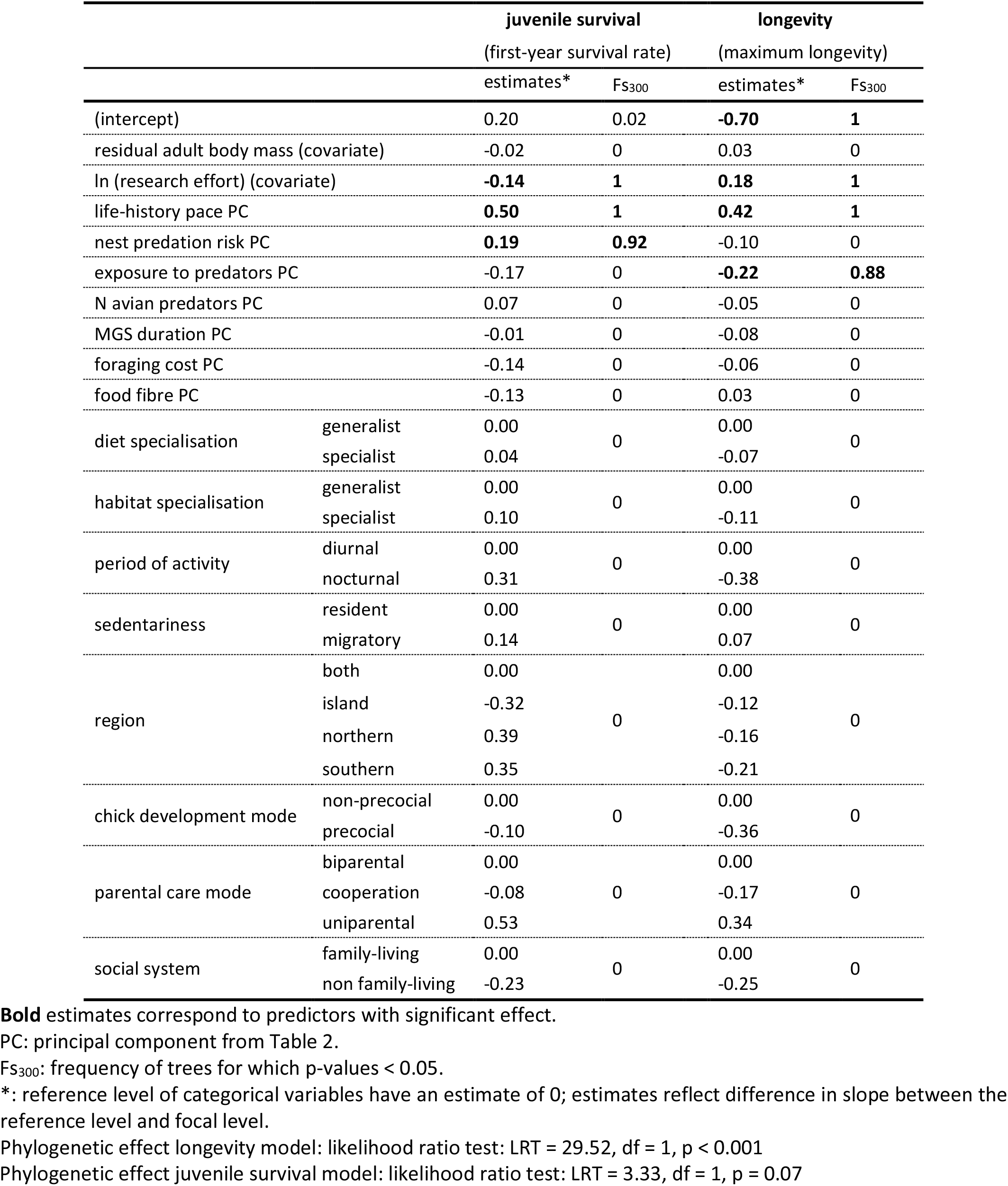
Correlates of juvenile survival and longevity. Results from phylogenetically controlled linear mixed-effect models testing the influence of key life-history, ecological and social traits on juvenile survival and longevity, respectively.

### Combinations of juvenile survival and longevity

Juvenile survival and longevity were positively correlated (r_Spearman_= 0.28, p <0.0001) (Figs. 1 and S5) and the slope of their linear regression was significant (N = 293, RMA slope = 53.15, 95% CI of the slope: 34.13, 81.71, p < 0.0001). However, there were major deviations from the regression line (R^2^ = 0.07), and 229 of 293 species (78%) fell outside the 95% confidence interval (CI) of the RMA regression (Figs. 1 and S6). We note that the percentage of species that deviate from the overall juvenile survival-longevity relationship was only a slightly lower (71%) when using a more conservative CI (99%CI: Fig. S7).

**Figure 1.**
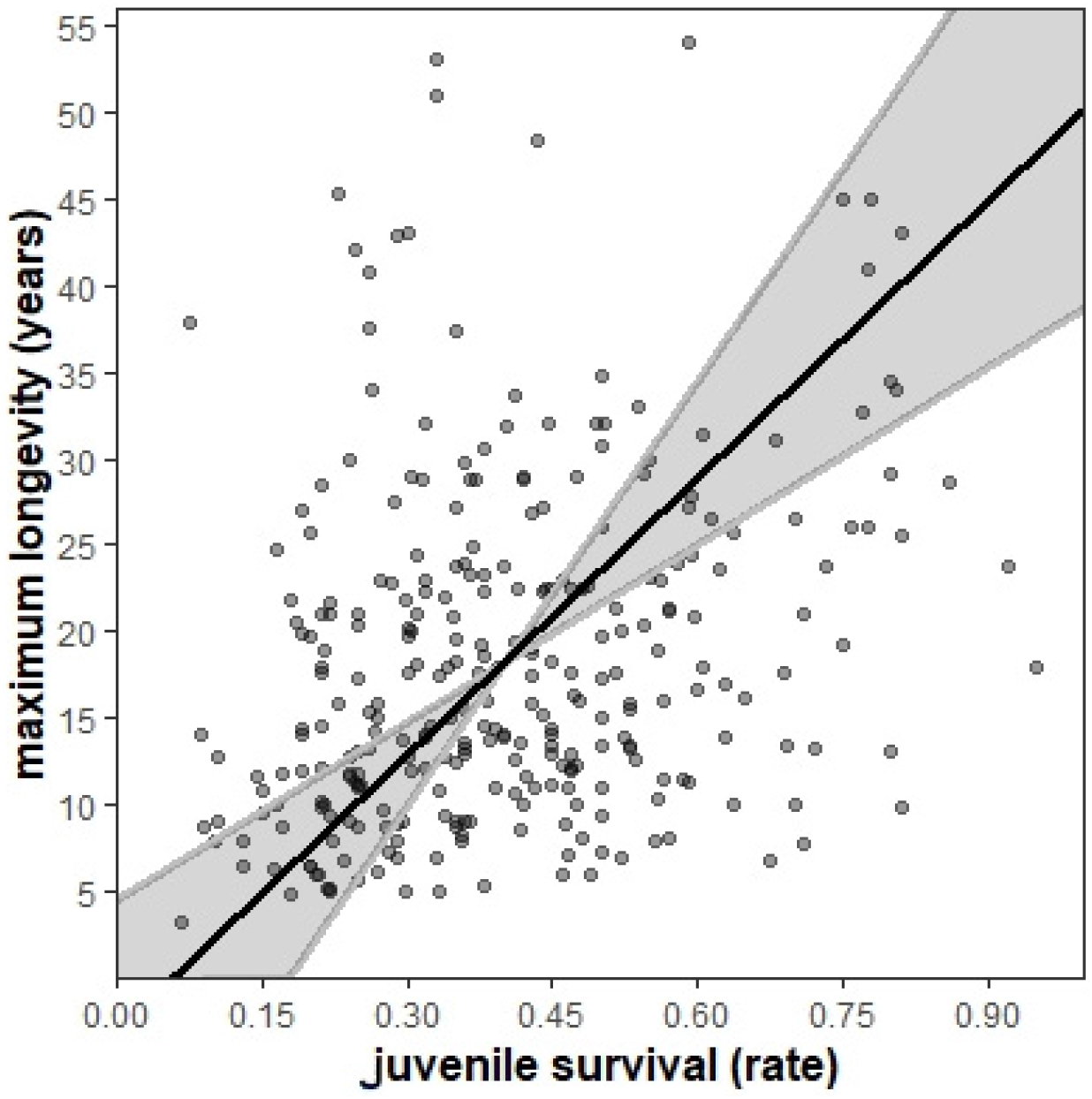
Correlation between juvenile survival (first-year survival) and maximum longevity on 293 bird species. RMA slope = 53.15, 95% CI (34.13, 81.71); r_spearman_ = 0.28, S = 3003600, p <0.0001. 64 species (22%) are inside and 229 (78%) outside the 95%CI of the regression line (shaded area). See Fig. S6 for species identification.

#### Positive associations between juvenile survival and longevity

In general, positive JS-L combinations were associated with family living, or a high risk of nest predation (open nest on or close to the ground, Table 1) but those effects were independent of the direction of the relationship (significant simple effects: Table S8). Opposite positive combinations of juvenile survival and longevity (low-low vs. high-high) were differently associated with specific-species attributes. This was reflected by the significant two-way interactions between the sign of PC1 (negative: low-low vs. positive: high-high JS-L combinations, Fig. 2) and sedentariness, exposure to predators, life-history pace and parental care mode. Species with high juvenile survival-high longevity combinations were migratory, had a low exposure to adult predators (pelagic forager, living in open habitat; Table 1), a slow life-history pace or uniparental care. In contrast, species with low juvenile survival-low longevity combinations were sedentary, had high exposure to predators, a fast life-history pace, or had bi-parental or cooperative offspring care) (Figs. 2 and S8, Table S8).

**Figure 2.**
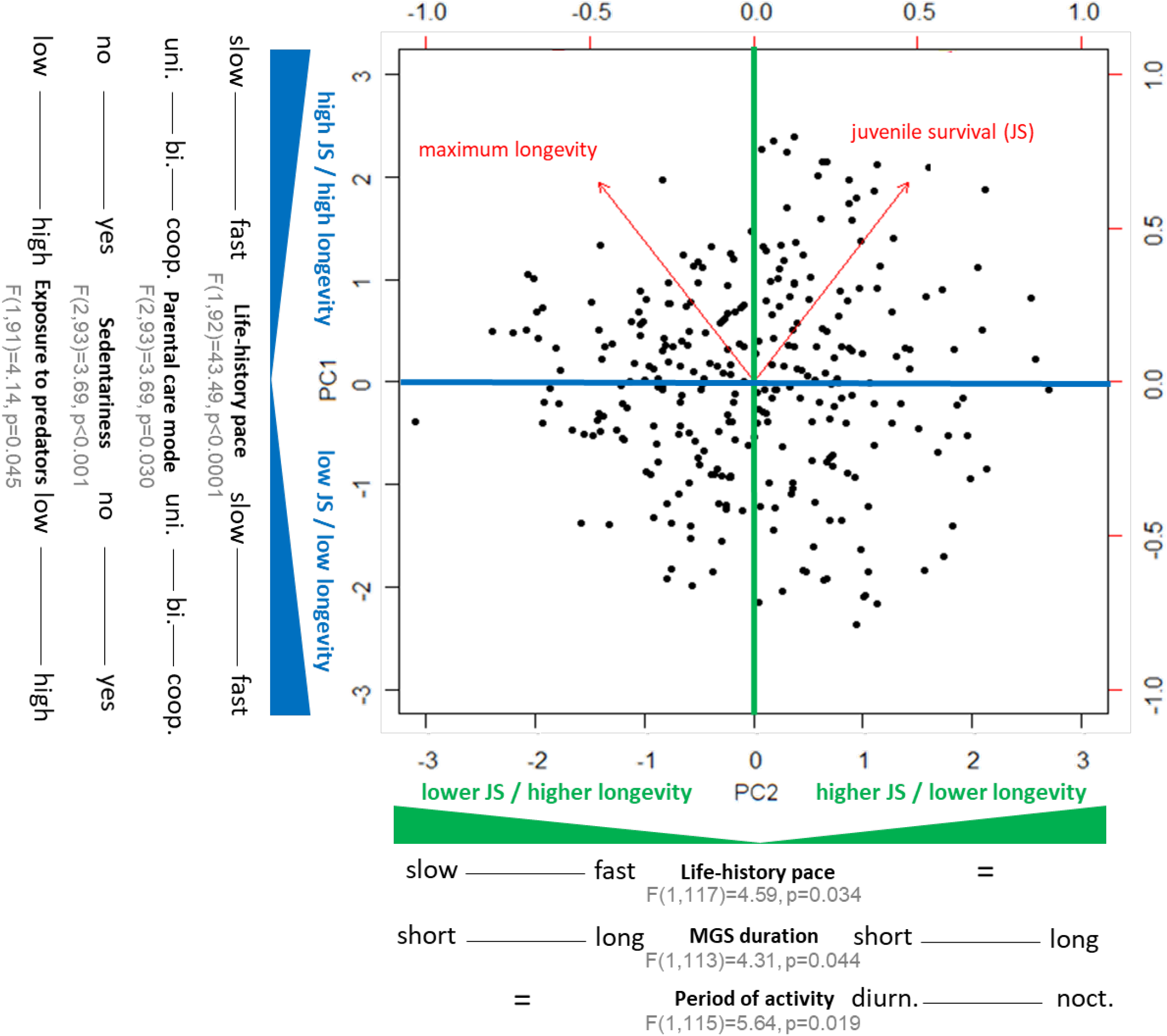
Correlates of the positive (PC1) and mismatching (PC2) combinations of juvenile survival and longevity. Graphical summary of the main results from the backward model selections on phylogenetically controlled linear mixed models investigating which life-history, ecological and social traits characterised species with different combinations of juvenile survival (first-year survival) and longevity (N=204). The blue axis (PC1) represents combinations that concur with tETA’s classical prediction (high juvenile survival associated with high longevity or vice versa). The green axis (PC2) represents combinations that deviate from tETA’s classical prediction (deviation towards higher juvenile survival associated with lower longevity or vice versa). Graphics of each independent results are provided in Figs. S8 and S9. See Fig. S10 for species identification and Fig. S11 for order identification. JS = juvenile survival, Coop. = cooperative breeding, bi. = biparental care, uni. = uniparental care, noct. = nocturnal, diurn. = diurnal, F(…,…) = Conditional F statistic and its degrees of freedoms averaged over the 300 models, p = averaged p value over the 300 models.

#### Mismatches between juvenile survival and longevity

In general, JS-L mismatches were associated with low exposure to adult predators (pelagic foraging, living in open habitat, Table 1) or being a habitat generalist, but these effects were independent of the direction of the relationship between juvenile survival and longevity (significant simple effects: Table S9). Opposite JS-L mismatches (low-high vs. high-low) were differently associated with specific-species attributes as reflected by the significant two-way interactions between the sign of PC2 (negative: low-high vs. positive: high-low JS-L combinations, Fig. 2) and period of activity, MGS duration and life-history pace. Species with stronger than expected combinations of high juvenile survival-low longevity lived in stable environments with long growing seasons (Table 1), or were nocturnal. In contrast, species with outstandingly low juvenile survival-high longevity combinations lived in variable environment with short growing seasons (Table 1) or had a slow life-history pace (Figs. 2 and S9, Table S9).

## Discussion

Empirical studies often use longevity as a proxy of life-history pace, based on the assumption of tETA that juvenile survival and longevity are positively correlated^2,8,12,94^. While this pattern is supported by previous work^2,6,7,12^ and is generally visible in our data, our analyses show that around 70% of bird species significantly deviate from this overall juvenile survival-longevity positive relationship (Figs. 1 and S6). Our analyses demonstrate that a wide range of survivals’ combinations evolved, some in accordance with the classical prediction of tETA while others contrasting it, partly supporting recent developments in this field^9,14–16^. Overall, this study raises awareness on the fact that the relationship between juvenile survival and longevity is not a black or white concept, but a range of grey nuances, and identifies key evolutionary forces driving the coevolution between juvenile survival and longevity.

### Correlates of juvenile survival and longevity

On average, a slow life-history pace (in our study corresponding to: large body size, low annual reproductive investment, long incubation period, Table 2) is associated with high juvenile survival and longevity (Table 3), supporting life-history theory^12,19^. However, while juvenile survival and longevity are positively correlated (Figs. 1 and S5), their individual variation are also associated with particular parameters (Table 3;^95^), supporting findings from mammals^21^. Our analyses show that nest predation risk (index based on nest location and nest type, Table 1) only influences juvenile survival while exposure to predators of adults (index of habitat openness, Table 1 and 2) only influences longevity (Table 3). Consequently, these factors are likely to play an important role in the evolution of diverse juvenile survival-longevity patterns.

Juveniles are often less conspicuous than adults due to more cryptic coloration and behaviours^96–99^, reducing their vulnerability to predation. Accordingly, a high exposure to predators of adults is associated with decreased longevity only (Table 3). In contrast, a low nest predation risk is associated with low juvenile survival only (Table 3). In this study, this latter association concerns mainly cavity-breeding species (Table 2) known to often experience a lower nest predation risk than open-nesting species^72^. However, in cavities, nestlings are often exposed to ectoparasites^100,101^, reducing their body condition^100,102,103^, potentially explaining a reduced juvenile survival in these species^95^ (Table 3). Therefore, nesting habits that provide short-term benefits early on in life may have negative down-streams effect on juvenile survival that so far were not anticipated (but see^42^).

### Combinations of juvenile survival and longevity

Most species (78%) deviate significantly from the positive juvenile survival-longevity regression revealing the existence of a continuum of patterns (Figs. 1 and S6), challenging the classical assumption of tETA^2,6,7,12^. The degree of this deviation varies considerably between species (Figs. 2 and S10), demonstrating that the association between juvenile survival and longevity evolved towards multiple adaptive combinations in birds. Some part of this mismatch may represent random variation and cannot be explained by consistent biological patterns. However, variation in survival at different life stages is likely to represent distinct strategies, shaped by natural selection to achieve the most optimal solutions in a given combination of external and internal factors. Thus, instead of forcing the long-accepted pattern of tETA or challenging it with opposing hypotheses, we should adopt a more diverse approach. Accordingly, one should embrace that various possible juvenile survival-longevity combinations exist (including the non-tETA compliant ones), and their actual values should be assumed to maximize population viability. Our framework integrating ecological, life-history and social moderators clearly demonstrates that such a heterogeneous picture is biologically more realistic.

Our analyses on the associations between juvenile survival and longevity do not allow us to investigate unusual juvenile survival and longevity separately, limiting our ability to identify underlying mechanisms. This would require an in-depth view of what is happening between individuals, calling for more comparative studies and experiments on both juvenile survival and longevity at the intra-species level. However, species-level deviations from the positive correlation between juvenile survival and longevity likely reflect that certain selective factors only influence specific life stages^35,104^. Patterns observed between different taxa can be thought of as averaged outcomes of selective pressures, acting over long periods of time. Indeed, age-dependent changes in body size, coloration, behaviour, or the onset of reproduction and senescence, can affect extrinsic and intrinsic mortality differently at different life stages^35,104^. For example, juveniles early on in life are often smaller than adults, making them more susceptible to predation^32,39^. Also, juvenile survival may be low in species that live in challenging environments, have elaborate foraging techniques or a specialised diet, as juveniles in those species seem to need more time to acquire adult skill levels^62,68,105^. In contrast, only adults pay costs of reproduction, which may reduce their longevity directly, or indirectly, for instance through increased exposure to predators as a consequence of increased foraging effort^106^, or displaying the own quality to potential partners^107^.

#### Positive associations between juvenile survival and longevity

Positive JS-L combinations are in accordance with the classical prediction of tETA, indicating that life-history, ecological, and social parameters have similar effects on juvenile survival and longevity. Our analyses show that high juvenile survival-high longevity combinations are found in species that are migratory, have a low exposure to predators, a slow life-history pace or uniparental care (Fig. 2), and are mostly observed in Accipitriformes, Anseriformes, Charadriiformes, and Pelicaniformes (Fig. S11). In contrast, low juvenile survival-low longevity combinations are found in species that are sedentary, have a high exposure to predators, a fast life-history pace, or have cooperative or biparental brood care (Fig. 2), and are mostly observed in Galliformes and Passeriformes (Fig. S11).

Migration is regularly found in species breeding at higher latitudes or altitudes, allowing them to escape harsh winter conditions^55^. In most of these species, juveniles and adults are migratory, thus affecting both life stages. While previous research showed that migration can be costly (i.e., being associated with smaller relative brain sizes;^108^), our results highlight that it has a positive effect on survival in general. Moreover, a low exposure to predators is beneficial for both juvenile and adults, making pelagic species particularly long-lived^32^. As predicted by life-history theory, species with a slow life-history pace have increased juvenile survival and longevity^12,19,66^. Furthermore, parental care is costly^54,109^. To ensure the survival of their offspring, parents provide them with food, thermoregulation, and protection from predators, which, on top of being energy demanding, exposes the parents to an increased risk of predation^54,110^. Thus, it seems surprising that species with uniparental care have combination of higher juvenile survival and longevity compared to biparental and cooperatively breeding species. A possible explanation is that particularly species with low costs of parental care evolved uniparental care, leading to increased juvenile survival and longevity. Clearly, this finding calls for further studies to investigate both the drivers and consequences of uniparental care.

#### Mismatches between juvenile survival and longevity

Mismatching combinations of juvenile survival and longevity suggest that certain factors specifically act upon juvenile survival or longevity, or have opposing effects on juvenile survival and longevity, leading to age-specific differences in survival. Our results demonstrate that high juvenile survival-low longevity combinations are found in species that live in stable environments with long growing seasons or are nocturnal (Fig. 2), and are mostly observed in Apodiformes and Galliformes (Fig. S11). In contrast, low juvenile survival-high longevity combinations are found in species that live in variable environments with short growing seasons or have a slow life-history pace (Fig. 2), and are mostly observed in Pelicaniformes and Procellariiformes (Fig. S11).

Conceivably, living in stable environments may particularly affect juvenile survival, reducing their winter mortality, while the opposite is the case in variable environments. The high juvenile survival-low longevity combinations found in nocturnal or crepuscular species is likely to reflect reduced juvenile mortality, given that most predators of birds are diurnal bird species^32^. In contrast, combinations of low juvenile survival-high longevity found in species with a slow life-history pace is likely to reflect that long-lived species particularly invest in longevity, at the expense of high juvenile survival in some species. Generally, interpreting those interactions is not straightforward. We urge further studies, especially longitudinal ones, to improve our understanding of the interesting interspecific patterns revealed here.

### Conclusions

Our comparative study provides novel insights into interspecific variation in juvenile survival, longevity and their combination in birds, and highlights the importance to consider agespecific survival to understand the evolution of life-history traits^22,25,26,42,111^. It increases our knowledge on the correlates of longevity and the under-studied juvenile survival and shows that most species deviate from the classical prediction of tETA. Our findings show that multiple adaptive combinations of juvenile survival and longevity evolved (more than commonly expected), some in accordance with tETA’s classical prediction while others contradict it. Accordingly, we call for a novel, more diverse, approach to understand the link between juvenile survival and longevity, and to move beyond the classical prediction of tETA. Our analyses demonstrate that positive JS-L combinations co-vary along the pace of life continuum, and JS-L mismatches co-vary with the length of the growing season, where long growing seasons promote juvenile survival, while short growing seasons promote longevity. Interestingly, sociality (parental care) only explains positive JS-L combinations, while ecological and life-history traits explain both positive JS-L combinations (sedentariness, exposure to predators, pace of life) and JS-L mismatches (length of growing season, period of activity, pace of life). Finally, our analysis emphasizes the need of not only studying typical patterns, predicted by accepted hypotheses – but also looking at outlying cases, that may embody genuine biological patterns rather than random deviations from assumed relationships.

Overall, this study reveals that the various combinations of juvenile survival and longevity observed are shaped by a distinct and limited set of species-specific life-history, ecological and social attributes. This may reflect divergent selection on each survival estimate, or that divergent agespecific survival is at the origin of diversity in species attributes^112^. Finally, species with unexpected age-specific survival relationships are more likely to evolve uncommon combination of life-history traits^28^. Thus, insights into key factors associating with unusual age-specific survival (such as the one found in this study) could contribute to a better understanding of life-history evolution^22,25–28,42,111^.

## Acknowledgements

We thank Carel van Schaik and Gretchen Wagner for discussions and comments on the manuscript. Kate Mears and Katharine Bowgen for the help with data compilation. This study was financed by the Swiss National Research Foundation (grant number PPOOP3_123520 and PPOOP3_150752, to MG), the Polish National Science Centre (Sonata Bis program, grant agreement no UMO-2015/18/E/NZ8/00505, to SMD) and the Australian Research Council (DECRA Fellowship number DE180100202).

## Authorship

M.G. and E.M. compiled part of the data, E.M. performed all statistical analyses and wrote the first draft of the manuscript. M.G. and S.M.D contributed suggestions and text to subsequent drafts. S.M.D helped with the statistical methods. All authors contributed to revisions and gave final approval for publication.

## Data accessibility

The datasets supporting this article have been uploaded as part of the Supporting Information and will be archived in Dryad. The data DOI will be included at the end of the article.

