## Supplementary information for "Increased juvenile survival may not be universally linked to longevity: ecological, social and life-history drivers of age-specific mortality in birds"

**This PDF file includes:**

Tables S1 to S9
Figures S1 to S11
SI Reference

**Table S1. Analysis with AFR:** Results of the Principal Component Analysis (PCA) with varimax rotation on the 12 continuous predictors on a subset of species for which AFR was available (N=188).

| | | | | **life-history pace** | **exposure to predators** | **food fibre** | **nest predation risk** | **N avian predators** | **foraging cost** | **MGS duration** | |
|---|---|---|---|---|---|---|---|---|---|---|---|
| category | transformation | variable | nº component | 1 | 7 | 2 | 6 | 3 | 4 | 5 | h2 |
| life-history | ln | adult body mass | | **0.90** | -0.22 | -0.12 | 0.15 | -0.02 | -0.20 | -0.11 | 0.95 |
| | none | annual parental investment | | **-0.90** | 0.09 | -0.15 | 0.12 | 0.11 | -0.03 | -0.01 | 0.88 |
| | sqrt | incubation period | | **0.77** | -0.31 | 0.22 | 0.01 | -0.08 | -0.31 | -0.03 | 0.84 |
| | none | nestling period | | 0.51 | 0.16 | 0.46 | -0.56 | -0.08 | -0.02 | 0.19 | 0.84 |
| | ln | age at first reproduction | | 0.60 | -0.36 | 0.23 | -0.16 | -0.25 | 0.31 | -0.15 | 0.75 |
| ecological | none | nest predation risk | | 0.04 | -0.12 | -0.13 | **0.93** | -0.13 | 0.10 | -0.07 | 0.92 |
| | none | foraging cost | | -0.16 | 0.04 | -0.13 | 0.11 | -0.03 | **0.93** | -0.06 | 0.93 |
| | none | calorie content of food | | 0.12 | 0.53 | 0.58 | 0.05 | 0.46 | -0.07 | -0.01 | 0.84 |
| | ln | fibre content of food | | -0.13 | 0.13 | **-0.90** | 0.22 | -0.05 | 0.13 | -0.09 | 0.92 |
| | none | foraging exposure | | -0.31 | **0.85** | -0.06 | -0.05 | 0.02 | -0.02 | 0.01 | 0.82 |
| | sqrt | vegetation cover | | -0.18 | **0.83** | 0.01 | -0.16 | -0.10 | 0.10 | 0.11 | 0.77 |
| | sqrt | N avian predators | | -0.20 | -0.07 | 0.08 | -0.12 | **0.92** | -0.03 | 0.01 | 0.91 |
| | none | MGS duration | | -0.08 | 0.09 | 0.08 | -0.11 | 0.01 | -0.06 | **0.97** | 0.99 |
| | | SS loadings | | 3.05 | 2.03 | 1.54 | 1.35 | 1.17 | 1.15 | 1.05 | |
| | | cumulative variance explained (%) | | 23 | 39 | 51 | 61 | 70 | 79 | 87 | |

We considered coefficients of correlation greater than 0.7 or less than -0.7 to be high loadings (highlighted in bold). h2 is the communality of the 7 components.

**Table S2. Analysis with AFR:** Results from phylogenetically controlled linear mixed-effect models testing the influence of key life-history, ecological and social traits on juvenile survival and longevity, respectively on a subset of species for which AFR was available (N=188).

| | | **juvenile survival** (first-year survival rate) | | **longevity** (maximum longevity) | |
|---|---|---|---|---|---|
| | | estimates* | $Fs_{300}$ | estimates * | $Fs_{300}$ |
| (intercept) | | 0.43 | 0.19 | **-0.90** | **1** |
| residual adult body mass (covariate) | | -0.02 | 0 | -0.03 | 0 |
| ln (research effort) (covariate) | | **-0.16** | **1** | **0.21** | **1** |
| life-history pace PC | | **0.49** | **1** | **0.44** | **1** |
| nest predation risk PC | | **0.21** | **1** | -0.12 | 0 |
| exposure to predators PC | | -0.15 | 0 | **-0.24** | **0.59** |
| N avian predators PC | | 0.08 | 0 | -0.05 | 0 |
| MGS duration PC | | -0.03 | 0 | -0.06 | 0 |
| foraging cost PC | | -0.12 | 0 | -0.09 | 0 |
| food fibre PC | | -0.14 | 0 | 0.06 | 0 |
| diet specialization | generalist | 0.00 | 0 | 0.00 | 0 |
| | specialist | 0.02 | | -0.05 | |
| habitat specialization | generalist | 0.00 | 0 | 0.00 | 0 |
| | specialist | 0.08 | | -0.08 | |
| period of activity | diurnal | 0.00 | 0 | 0.00 | 0 |
| | nocturnal | 0.14 | | -0.37 | |
| sedentariness | resident | 0.00 | 0 | 0.00 | 0 |
| | migratory | 0.16 | | 0.04 | |
| region | both | 0.00 | 0 | 0.00 | 0 |
| | island | -0.23 | | -0.25 | |
| | northern | 0.34 | | -0.11 | |
| | southern | 0.45 | | -0.07 | |
| chick development mode | non-precocial | 0.00 | 0 | 0.00 | 0 |
| | precocial | -0.15 | | -0.28 | |
| parental care mode | biparental | 0.00 | 0 | 0.00 | 0 |
| | cooperation | -0.17 | | -0.20 | |
| | uniparental | 0.65 | | 0.31 | |
| social system | family-living | 0.00 | 0 | 0.00 | 0 |
| | non family-living | -0.27 | | -0.21 | |

**Bold** estimates correspond to predictors with significant effect.
PC: principal component from Table S1.
$Fs_{300}$: frequency of trees for which p-values < 0.05.
*: reference level of categorical variables have an estimate of 0; estimates reflect difference in slope between the reference level and focal level.
Phylogenetic effect longevity model: likelihood ratio test: LRT = 24.43, df = 1, p < 0.001
Phylogenetic effect juvenile survival model: likelihood ratio test: LRT = 2.67, df = 1, p = 0.10

**Table S3. Analysis with AFR:** Results from backward model selection on phylogenetically controlled linear mixed-effect model investigating which life-history, ecological and social traits characterize species with positive JS-L combinations (PC1) on a subset of species for which AFR was available (N=188).

| predictors | | estimates* | 95% CI | $Fs_{300}$ | P |
|---|---|---|---|---|---|
| *simples effects:* | | | | | |
| **(intercept)** | | **1.67** | **(1.67,1.67)** | **1** | **<0.0001** |
| residual adult body mass (covariate) | | -0.07 | (-0.07,-0.07) | 0 | 0.320 |
| **ln (research effort) (covariate)** | | **-0.10** | **(-0.10,-0.10)** | **1** | **0.002** |
| life-history pace | | -0.26 | (-0.26,-0.26) | 0 | 0.287 |
| **nest predation risk** | | **0.14** | **(0.14,0.14)** | **1** | **0.002** |
| exposure to predators | | 0.03 | (0.03,0.03) | 0 | 0.399 |
| sedentariness | resident | 0.00 | na | 0 | 0.264 |
| | migratory | -0.32 | (-0.32,-0.32) | | |
| social system | family-living | 0.00 | na | 0 | 0.073 |
| | non family-living | -0.20 | (-0.20,-0.20) | | |
| **parental care mode** | **biparental** | **0.00** | **na** | **0.67** | **0.044** |
| | **cooperation** | **-0.48** | **(-0.48,-0.48)** | | |
| | **uniparental** | **-0.63** | **(-0.63,-0.62)** | | |
| sign PC1 | negative | 0.00 | na | 0 | 0.338 |
| | positive | 0.35 | (0.35,0.35) | | |
| *Interactions:* | | | | | |
| **sign PC1 : life-history pace** | **negative** | **0.00** | **na** | **1** | **<0.0001** |
| | **positive** | **0.55** | **(0.55,0.55)** | | |
| sign PC1 : exposure to predators | negative | 0.00 | na | 0 | 0.085 |
| | positive | -0.14 | (-0.14,-0.14) | | |
| **sign PC1 : sedentariness** | **negative : resident** | **0.00** | **na** | **1** | **0.006** |
| | **negative : migratory** | **0.00** | **na** | | |
| | **positive : resident** | **0.00** | **na** | | |
| | **positive : migratory** | **0.49** | **(0.49,0.49)** | | |
| **sign PC1 : parental care mode** | **negative : biparental** | **0.00** | **na** | **1** | **0.009** |
| | **negative : cooperation** | **0.00** | **na** | | |
| | **negative : uniparental** | **0.00** | **na** | | |
| | **positive : biparental** | **0.00** | **na** | | |
| | **positive : cooperation** | **0.49** | **(0.49,0.49)** | | |
| | **positive : uniparental** | **0.81** | **(0.81,0.81)** | | |

**Bold** estimates correspond to predictors with significant effect. na – not applicable.

95% CI: confidence interval of the average estimate on the 300 trees, narrow CI means that the estimate is robust.

$Fs_{300}$: frequency of trees for which p-values < 0.05.

P: average p-value on the 300 trees.

*: reference level of categorical variables have an estimate of 0; estimates reflect difference in slope between the reference level and focal level.

Negative sign of PC1 refers to low juvenile survival-low longevity combinations; Positive sign of PC1 refers to high juvenile survival-high longevity combinations.

Phylogenetic effect: likelihood ratio test: LRT = 1.89, df = 1, p = 0.17

**Table S4. Analysis with AFR:** Results from backward model selection on phylogenetically controlled linear mixed-effect model investigating which life-history, ecological and social traits characterize species with JS-L mismatches (PC2) on a subset of species for which AFR was available (N=188).

| predictors | | estimates* | 95% CI | $Fs_{300}$ | P |
|---|---|---|---|---|---|
| *simples effects:* | | | | | |
| **(intercept)** | | **1.29** | **(1.29,1.29)** | **1** | **<0.0001** |
| residual adult body mass (covariate) | | 0.01 | (0.01,0.01) | 0 | 0.899 |
| **ln (research effort) (covariate)** | | **-0.08** | **(-0.08,-0.08)** | **1** | **0.034** |
| **life-history pace** | | **0.23** | **(0.23,0.23)** | **0.7** | **0.048** |
| MGS duration | | -0.11 | (-0.11,-0.11) | 0 | 0.711 |
| **exposure to predators** | | **-0.18** | **(-0.18,-0.18)** | **1** | **0.009** |
| sign PC2 | negative | 0.00 | na | 0 | 0.246 |
| | positive | -0.16 | (-0.16,-0.16) | | |
| period of activity | diurnal | 0.00 | na | 0 | 0.125 |
| | nocturnal | 0.06 | (0.06,0.06) | | |
| *Interactions:* | | | | | |
| **sign PC2 : life-history pace** | **negative** | **0.00** | **na** | **1** | **0.009** |
| | **positive** | **-0.21** | **(-0.21,-0.21)** | | |
| **sign PC2 : MGS duration** | **negative** | **0.00** | **na** | **1** | **0.024** |
| | **positive** | **0.18** | **(0.18,0.18)** | | |
| **sign PC2 : period of activity** | **negative : diurnal** | **0.00** | **na** | **1** | **0.024** |
| | **negative : nocturnal** | **0.00** | **na** | | |
| | **positive : diurnal** | **0.00** | **na** | | |
| | **positive : nocturnal** | **0.75** | **(0.75,0.75)** | | |

**Bold** estimates correspond to predictors with significant effect. na – not applicable.
95% CI: confidence interval of the average estimate on the 300 trees, narrow CI means that the estimate is robust.
$Fs_{300}$: frequency of trees for which p-values < 0.05.
P: average p-value on the 300 trees.
*: reference level of categorical variables have an estimate of 0; estimates reflect difference in slope between the reference level and focal level.
Negative sign of PC2 refers to low juvenile survival-high longevity combinations; Positive sign of PC1 refers to high juvenile survival-low longevity combinations.
Phylogenetic effect: likelihood ratio test: LRT = 5.85, df = 1, p =0.02

**Table S5. Correlation matrix**, standardized principal components loadings, and communality (h2) of juvenile survival and maximum longevity.

| correlation matrix | | | loadings | | $h^2$ |
|---|---|---|---|---|---|
| | ln(juvenile survival) | ln(maximum longevity) | PC1 | PC2 | |
| ln(juvenile survival) | 1 | 0.27 | 0.81 | -0.59 | 1 |
| ln(maximum longevity) | 0.27 | 1 | 0.81 | 0.59 | 0.78 |
| eigenvalue | | | 1.31 | 0.69 | |
| cumulative variance explained (%) | | | 65 | 100 | |

**Table S6.** Results from the full mixed model on absolute PC1 (positive JS-L combinations) (N=204).

| predictors | | estimates* | 95% CI | $Fs_{300}$ |
|---|---|---|---|---|
| *simples effects:* | | | | |
| **(intercept)** | | **1.64** | **(1.64,1.64)** | **1** |
| residual adult body mass (covariate) | | -0.10 | (-0.10,-0.10) | 0 |
| **ln (research effort) (covariate)** | | **-0.10** | **(-0.10,-0.10)** | **1** |
| life-history pace | | -0.25 | (-0.25,-0.25) | 0 |
| nest predation risk | | 0.05 | (0.05,0.05) | 0 |
| exposure to predators | | 0.07 | (0.07,0.07) | 0 |
| N avian predators | | -0.08 | (-0.08,-0.08) | 0 |
| MGS duration | | -0.04 | (-0.04,-0.04) | 0 |
| foraging cost | | -0.01 | (-0.01,-0.01) | 0 |
| food fibre | | 0.11 | (0.11,0.11) | 0 |
| diet specialization | generalist | 0.00 | na | 0 |
| | specialist | 0.03 | (0.03,0.03) | |
| habitat specialization | generalist | 0.00 | na | 0 |
| | specialist | -0.00 | (-0.00,-0.00) | |
| period of activity | diurnal | 0.00 | na | 0 |
| | nocturnal | -0.21 | (-0.21,-0.21) | |
| sedentariness | resident | 0.00 | na | 0 |
| | migratory | -0.24 | (-0.25,-0.24) | |
| region | both | 0.00 | na | 0 |
| | island | -0.05 | (-0.05,-0.05) | |
| | northern | -0.15 | (-0.15,-0.15) | |
| | southern | -0.11 | (-0.11,-0.11) | |
| chick development mode | non-precocial | 0.00 | na | 0 |
| | precocial | 0.53 | (0.53,0.53) | |
| **parental care mode** | **biparental** | **0.00** | **na** | **0.60** |
| | **cooperation** | **-0.30** | **(-0.30,-0.30)** | |
| | **uniparental** | **-0.88** | **(-0.88,-0.88)** | |
| social system | family-living | 0.00 | na | 0 |
| | non family-living | -0.11 | (-0.12,-0.11) | |
| sign PC1 | negative | 0.00 | na | 0 |
| | positive | -0.55 | (-0.55,-0.54) | |
| *Interactions:* | | | | |
| **sign PC1 : life-history pace** | **negative** | **0.00** | **na** | **1** |
| | **positive** | **0.53** | **(0.53,0.53)** | |
| sign PC1 : nest predation risk | negative | 0.00 | na | 0 |
| | positive | 0.03 | (0.03,0.03) | |
| **sign PC1 : exposure to predators** | **negative** | **0.00** | **na** | **1** |
| | **positive** | **-0.28** | **(-0.28,-0.28)** | |
| sign PC1 : N avian predators | negative | 0.00 | na | 0 |
| | positive | 0.10 | (0.10,0.10) | |
| sign PC1 : MGS duration | negative | 0.00 | na | 0 |
| | positive | 0.11 | (0.11,0.11) | |
| sign PC1 : foraging cost | negative | 0.00 | na | 0 |
| | positive | -0.13 | (-0.13,-0.13) | |
| sign PC1 : food fibre | negative | 0.00 | na | 0 |
| | positive | -0.06 | (-0.06,-0.06) | |

**Table S6 following.** Results from the full mixed model on absolute PC1 (positive JS-L combinations) (N=204).

| predictors | | estimates* | 95% CI | $Fs_{300}$ |
|---|---|---|---|---|
| *Interactions:* | | | | |
| sign PC1 : diet specialization | negative : generalist | 0.00 | na | 0 |
| | negative : specialist | 0.00 | na | |
| | positive : generalist | 0.00 | na | |
| | positive : specialist | -0.18 | (-0.18,-0.17) | |
| sign PC1 : habitat specialization | negative : generalist | 0.00 | na | 0 |
| | negative : specialist | 0.00 | na | |
| | positive : generalist | 0.00 | na | |
| | positive : specialist | -0.02 | (-0.02,-0.02) | |
| sign PC1 : period of activity | negative : diurnal | 0.00 | na | 0 |
| | negative : nocturnal | 0.00 | na | |
| | positive : diurnal | 0.00 | na | |
| | positive : nocturnal | 0.05 | (0.05,0.05) | |
| **sign PC1 : sedentariness** | **negative : resident** | **0.00** | **na** | **1** |
| | **negative : migratory** | **0.00** | **na** | |
| | **positive : resident** | **0.00** | **na** | |
| | **positive : migratory** | **0.52** | **(0.52,0.52)** | |
| sign PC1 : region | negative : both | 0.00 | na | 0 |
| | negative : island | 0.00 | na | |
| | negative : northern | 0.00 | na | |
| | negative : southern | 0.00 | na | |
| | positive : both | 0.00 | na | |
| | positive : island | 0.09 | (0.09,0.09) | |
| | positive : northern | 0.43 | (0.43,0.43) | |
| | positive : southern | 0.03 | (0.03,0.03) | |
| sign PC1 : chick development mode | negative : non-precocial | 0.00 | na | 0 |
| | negative : precocial | 0.00 | na | |
| | positive : non-precocial | 0.00 | na | |
| | positive : precocial | -0.51 | (-0.51,-0.51) | |
| sign PC1 : parental care mode | negative : biparental | 0.00 | na | 0 |
| | negative : cooperation | 0.00 | na | |
| | negative : uniparental | 0.00 | na | |
| | positive : biparental | 0.00 | na | |
| | positive : cooperation | 0.33 | (0.33,0.33) | |
| | positive : uniparental | 0.73 | (0.73,0.73) | |
| sign PC1 : social system | negative : family-living | 0.00 | na | 0 |
| | negative : non family-living | 0.00 | na | |
| | positive : family-living | 0.00 | na | |
| | positive : non family-living | -0.18 | (-0.18,-0.18) | |

**Bold** estimates correspond to predictors with significant effect. na – not applicable.
95% CI: confidence interval of the average estimate on the 300 trees, narrow CI means that the estimate is robust.
$Fs_{300}$: frequency of trees for which p-values < 0.05.
*: reference level of categorical variables have an estimate of 0; estimates reflect difference in slope between the reference level and focal level.
Negative sign of PC1 refers to low juvenile survival-low longevity combinations; Positive sign of PC1 refers to high juvenile survival-high longevity combinations.
Phylogenetic effect: likelihood ratio test: LRT = 3.25, df = 1, p = 0.07

**Table S7.** Results from the full mixed model on absolute PC2 (JS-L mismatches) (N=204).

| predictors | | estimates* | 95% CI | $Fs_{300}$ |
|---|---|---|---|---|
| *simples effects:* | | | | |
| (intercept) | | 1.36 | (1.35,1.36) | 1 |
| residual adult body mass (covariate) | | -0.00 | (-0.00,-0.00) | 0 |
| ln (research effort) (covariate) | | -0.05 | (-0.05,-0.05) | 0 |
| life-history pace | | 0.21 | (0.21,0.21) | 0 |
| nest predation risk | | -0.04 | (-0.04,-0.04) | 0 |
| **exposure to predators** | | **-0.22** | **(-0.22,-0.22)** | **1** |
| N avian predators | | -0.00 | (-0.00,-0.00) | 0 |
| MGS duration | | -0.10 | (-0.11,-0.10) | 0 |
| foraging cost | | 0.00 | (0.00,0.00) | 0 |
| food fibre | | 0.02 | (0.02,0.02) | 0 |
| diet specialization | generalist | 0.00 | na | 0 |
| | specialist | -0.03 | (-0.03,-0.02) | |
| **habitat specialization** | **generalist** | **0.00** | **na** | **1** |
| | **specialist** | **-0.22** | **(-0.22,-0.22)** | |
| period of activity | diurnal | 0.00 | na | 0 |
| | nocturnal | -0.07 | (-0.07,-0.07) | |
| sedentariness | resident | 0.00 | na | 0 |
| | migratory | -0.12 | (-0.12,-0.12) | |
| region | both | 0.00 | na | 0 |
| | island | 0.15 | (0.15,0.15) | |
| | northern | -0.06 | (-0.06,-0.06) | |
| | southern | -0.10 | (-0.10,-0.10) | |
| chick development mode | non-precocial | 0.00 | na | 0 |
| | precocial | -0.13 | (-0.13,-0.13) | |
| parental care mode | biparental | 0.00 | na | 0 |
| | cooperation | 0.05 | (0.05,0.05) | |
| | uniparental | 0.40 | (0.40,0.40) | |
| social system | family-living | 0.00 | na | 0 |
| | non family-living | -0.05 | (-0.05,-0.05) | |
| sign PC2 | negative | 0.00 | na | 0 |
| | positive | -0.38 | (-0.39,-0.38) | |
| *Interactions:* | | | | |
| sign PC2 : life-history pace | negative | 0.00 | na | 0 |
| | positive | -0.17 | (-0.17,-0.17) | |
| sign PC2 : nest predation risk | negative | 0.00 | na | 0 |
| | positive | 0.05 | (0.05,0.05) | |
| sign PC2 : exposure to predators | negative | 0.00 | na | 0 |
| | positive | -0.02 | (-0.02,-0.02) | |
| sign PC2 : N avian predators | negative | 0.00 | na | 0 |
| | positive | -0.00 | (-0.00,-0.00) | |
| sign PC2 : MGS duration | negative | 0.00 | na | 0 |
| | positive | 0.18 | (0.17,0.18) | |
| sign PC2 : foraging cost | negative | 0.00 | na | 0 |
| | positive | 0.02 | (0.02,0.02) | |
| sign PC2 : food fibre | negative | 0.00 | na | 0 |
| | positive | -0.03 | (-0.03,-0.03) | |

**Table S7 following.** Results from the full mixed model on absolute PC2 (JS-L mismatches) (N=204).

| predictors | | estimates* | 95% CI | $Fs_{300}$ |
|---|---|---|---|---|
| *Interactions:* | | | | |
| sign PC2 : diet specialization | negative : generalist | 0.00 | na | 0 |
| | negative : specialist | 0.00 | na | |
| | positive : generalist | 0.00 | na | |
| | positive : specialist | 0.03 | (0.03,0.03) | |
| sign PC2 : habitat specialization | negative : generalist | 0.00 | na | 0 |
| | negative : specialist | 0.00 | na | |
| | positive : generalist | 0.00 | na | |
| | positive : specialist | 0.03 | (0.03,0.03) | |
| **sign PC2 : period of activity** | **negative : diurnal** | **0.00** | **na** | **1** |
| | **negative : nocturnal** | **0.00** | **na** | |
| | **positive : diurnal** | **0.00** | **na** | |
| | **positive : nocturnal** | **0.84** | **(0.84,0.84)** | |
| sign PC2 : sedentariness | negative : resident | 0.00 | na | 0 |
| | negative : migratory | 0.00 | na | |
| | positive : resident | 0.00 | na | |
| | positive : migratory | -0.06 | (-0.06,-0.06) | |
| sign PC2 : region | negative : both | 0.00 | na | 0 |
| | negative : island | 0.00 | na | |
| | negative : northern | 0.00 | na | |
| | negative : southern | 0.00 | na | |
| | positive : both | 0.00 | na | |
| | positive : island | 0.42 | (0.42,0.42) | |
| | positive : northern | 0.27 | (0.27,0.27) | |
| | positive : southern | 0.09 | (0.09,0.09) | |
| sign PC2 : chick development mode | negative : non-precocial | 0.00 | na | 0 |
| | negative : precocial | 0.00 | na | |
| | positive : non-precocial | 0.00 | na | |
| | positive : precocial | 0.24 | (0.24,0.25) | |
| sign PC2 : parental care mode | negative : biparental | 0.00 | na | 0 |
| | negative : cooperation | 0.00 | na | |
| | negative : uniparental | 0.00 | na | |
| | positive : biparental | 0.00 | na | |
| | positive : cooperation | 0.19 | (0.19,0.19) | |
| | positive : uniparental | -0.28 | (-0.28,-0.28) | |
| sign PC2 : social system | negative : family-living | 0.00 | na | 0 |
| | negative : non family-living | 0.00 | na | |
| | positive : family-living | 0.00 | na | |
| | positive : non family-living | 0.03 | (0.03,0.03) | |

**Bold** estimates correspond to predictors with significant effect. na – not applicable.
95% CI: confidence interval of the average estimate on the 300 trees, narrow CI means that the estimate is robust.
$Fs_{300}$: frequency of trees for which p-values < 0.05.
*: reference level of categorical variables have an estimate of 0; estimates reflect difference in slope between the reference level and focal level.
Negative sign of PC2 refers to low juvenile survival-high longevity combinations; Positive sign of PC1 refers to high juvenile survival-low longevity combinations.
Phylogenetic effect: likelihood ratio test: LRT = 0.86, df = 1, p = 0.35

**Table S8.** Results from backward model selection on phylogenetically controlled linear mixed-effect model investigating which life-history, ecological and social traits characterize species with positive JS-L combinations (PC1) (N=204).

| predictors | | estimates* | 95% CI | $Fs_{300}$ | P |
|---|---|---|---|---|---|
| *simples effects:* | | | | | |
| **(intercept)** | | **1.51** | **(1.50,1.51)** | **1** | **<0.0001** |
| residual adult body mass (covariate) | | -0.06 | (-0.06,-0.06) | 0 | 0.359 |
| **ln (research effort) (covariate)** | | **-0.07** | **(-0.07,-0.07)** | **1** | **0.016** |
| life-history pace | | -0.30 | (-0.30,-0.30) | 0 | 0.736 |
| **nest predation risk** | | **0.12** | **(0.12,0.12)** | **1** | **0.004** |
| exposure to predators | | 0.05 | (0.05,0.05) | 0 | 0.414 |
| sedentariness | resident | 0.00 | na | 0 | 0.664 |
| | migratory | -0.29 | (-0.29,-0.29) | | |
| **social system** | **family-living** | **0.00** | **na** | **1** | **0.017** |
| | **non family-living** | **-0.25** | **(-0.25,-0.25)** | | |
| **parental care mode** | **biparental** | **0.00** | **na** | **0.70** | **0.042** |
| | **cooperation** | **-0.44** | **(-0.44,-0.43)** | | |
| | **uniparental** | **-0.56** | **(-0.57,-0.56)** | | |
| sign PC1 | negative | 0.00 | na | 0 | 0.231 |
| | positive | -0.53 | (-0.53,-0.53) | | |
| *Interactions:* | | | | | |
| **sign PC1 : life-history pace** | **negative** | **0.00** | **na** | **1** | **<0.0001** |
| | **positive** | **0.55** | **(0.55,0.55)** | | |
| **sign PC1 : exposure to predators** | **negative** | **0.00** | **na** | **0.88** | **0.045** |
| | **positive** | **-0.15** | **(-0.15,-0.15)** | | |
| **sign PC1 : sedentariness** | **negative : resident** | **0.00** | **na** | **1** | **<0.001** |
| | **negative : migratory** | **0.00** | **na** | | |
| | **positive : resident** | **0.00** | **na** | | |
| | **positive : migratory** | **0.56** | **(0.55,0.56)** | | |
| **sign PC1 : parental care mode** | **negative : biparental** | **0.00** | **na** | **0.97** | **0.029** |
| | **negative : cooperation** | **0.00** | **na** | | |
| | **negative : uniparental** | **0.00** | **na** | | |
| | **positive : biparental** | **0.00** | **na** | | |
| | **positive : cooperation** | **0.39** | **(0.39,0.39)** | | |
| | **positive : uniparental** | **0.63** | **(0.63,0.64)** | | |

**Bold** estimates correspond to predictors with significant effect. na – not applicable.

95% CI: confidence interval of the average estimate on the 300 trees, narrow CI means that the estimate is robust.

$Fs_{300}$: frequency of trees for which p-values < 0.05.

P: average p-value on the 300 trees.

*: reference level of categorical variables have an estimate of 0; estimates reflect difference in slope between the reference level and focal level.

Negative sign of PC1 refers to low juvenile survival-low longevity combinations; Positive sign of PC1 refers to high juvenile survival-high longevity combinations.

Phylogenetic effect: likelihood ratio test: LRT = 0.10, df = 1, p = 0.75

**Table S9.** Results from backward model selection on phylogenetically controlled linear mixed-effect model investigating which life-history, ecological and social traits characterize species with JS-L mismatches (PC2) (N=204).

| predictors | | estimates* | 95% CI | $Fs_{300}$ | P |
|---|---|---|---|---|---|
| *simples effects:* | | | | | |
| **(intercept)** | | **1.34** | **(1.34,1.34)** | **1** | **<0.0001** |
| residual adult body mass (covariate) | | 0.01 | (0.01,0.01) | 0 | 0.860 |
| ln (research effort) (covariate) | | -0.06 | (-0.06,-0.06) | 0 | 0.080 |
| **life-history pace** | | **0.21** | **(0.21,0.21)** | **1** | **0.026** |
| MGS duration | | -0.10 | (-0.10,-0.10) | 0 | 0.531 |
| **exposure to predators** | | **-0.21** | **(-0.21,-0.21)** | **1** | **<0.001** |
| sign PC2 | negative | 0.00 | na | 0 | 0.426 |
| | positive | -0.12 | (-0.12,-0.12) | | |
| period of activity | diurnal | 0.00 | na | 0 | 0.064 |
| | nocturnal | 0.02 | (0.02,0.02) | | |
| **habitat specialization** | **generalist** | **0.00** | **na** | **1** | **0.044** |
| | **specialist** | **-0.17** | **(-0.17,-0.17)** | | |
| sedentariness | resident | 0.00 | na | 0 | 0.086 |
| | migratory | -0.16 | (-0.16,-0.16) | | |
| *Interactions:* | | | | | |
| **sign PC2 : life-history pace** | **negative** | **0.00** | **na** | **1** | **0.034** |
| | **positive** | **-0.16** | **(-0.16,-0.16)** | | |
| **sign PC2 : MGS duration** | **negative** | **0.00** | **na** | **1** | **0.040** |
| | **positive** | **0.15** | **(0.15,0.16)** | | |
| **sign PC2 : period of activity** | **negative : diurnal** | **0.00** | **na** | **1** | **0.019** |
| | **negative : nocturnal** | **0.00** | **na** | | |
| | **positive : diurnal** | **0.00** | **na** | | |
| | **positive : nocturnal** | **0.68** | **(0.68,0.68)** | | |

**Bold** estimates correspond to predictors with significant effect. na – not applicable.
95% CI: confidence interval of the average estimate on the 300 trees, narrow CI means that the estimate is robust.
$Fs_{300}$: frequency of trees for which p-values < 0.05.
P: average p-value on the 300 trees.
*: reference level of categorical variables have an estimate of 0; estimates reflect difference in slope between the reference level and focal level.
Negative sign of PC2 refers to low juvenile survival-high longevity combinations; Positive sign of PC1 refers to high juvenile survival-low longevity combinations.
Phylogenetic effect: likelihood ratio test: LRT = 3.25, df = 1, p =0.07

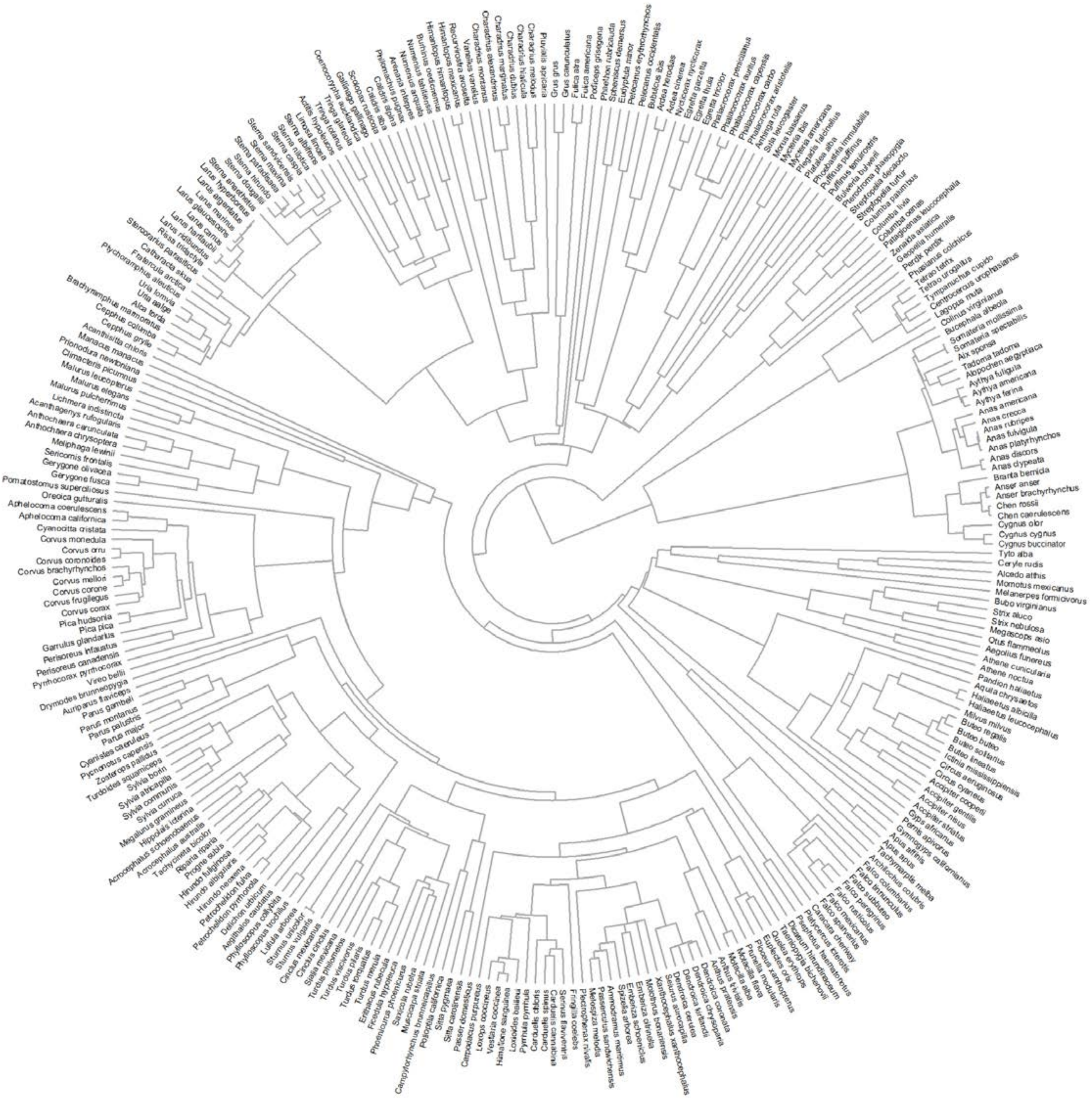


**Fig. S1.** Phylogenetic tree for the set of 293 species (based on the full tree from 1).

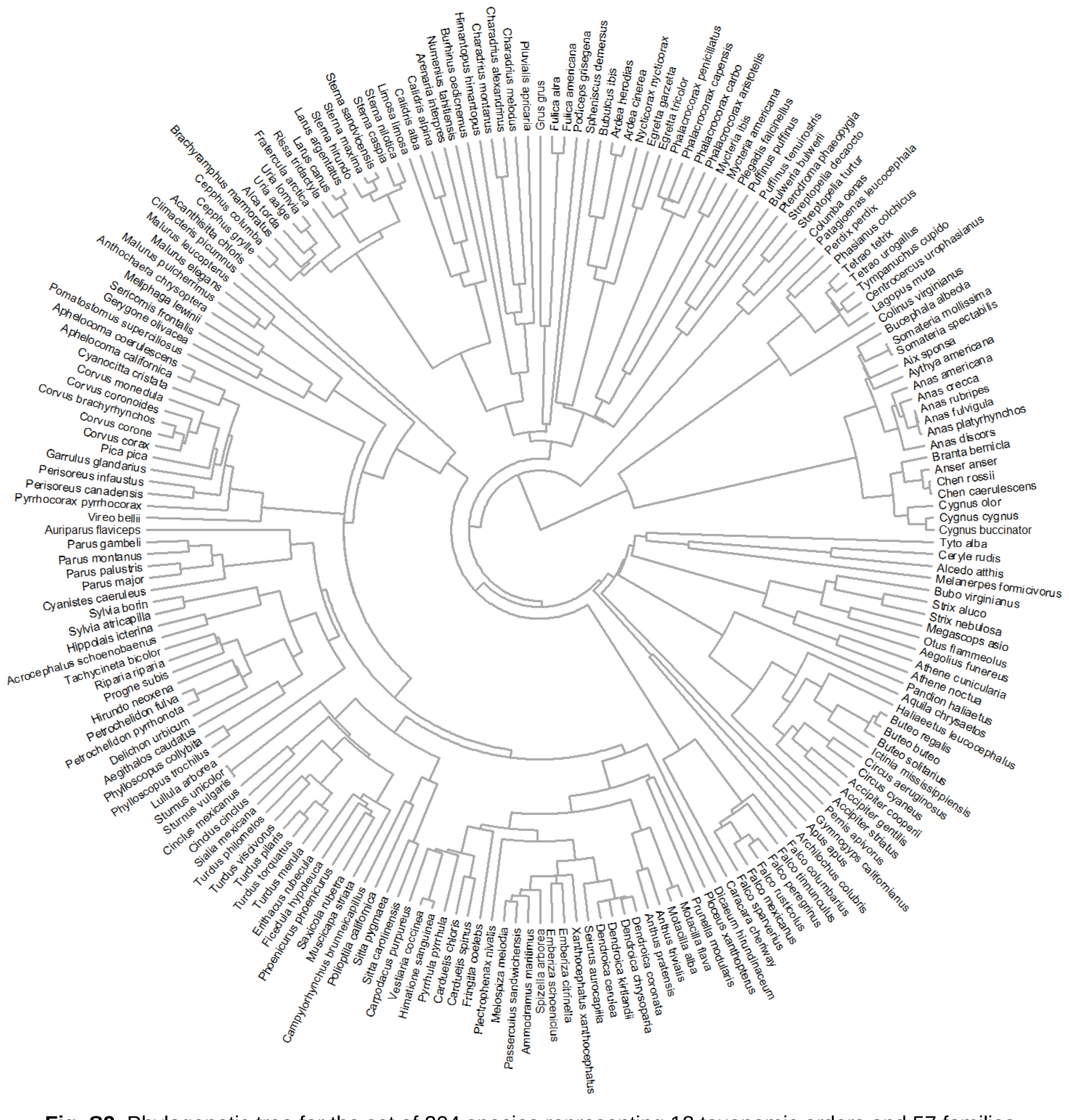


**Fig. S2.** Phylogenetic tree for the set of 204 species representing 18 taxonomic orders and 57 families (based on the full tree from 1).

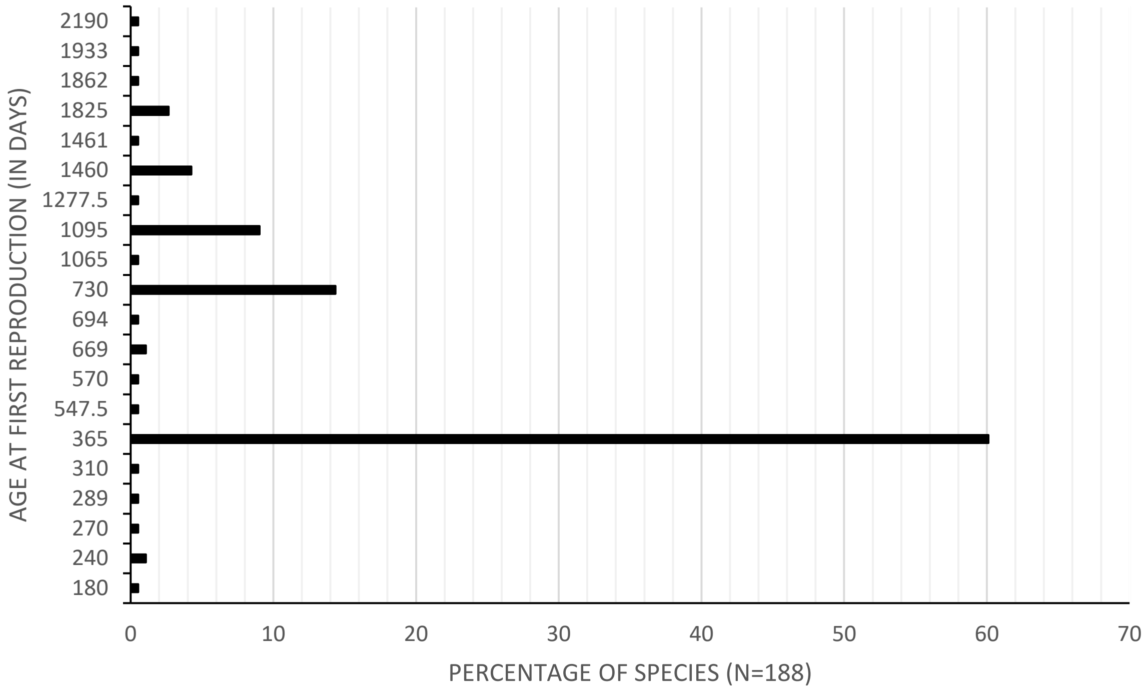


**Fig. S3.** Frequency of observation of the different age at first reproduction on 188 species (63.8% of the species had an AFR ≤ 1 year old, 17% ]1; 2], 9.6% ]2; 3] and 9.6% > 3 years old).

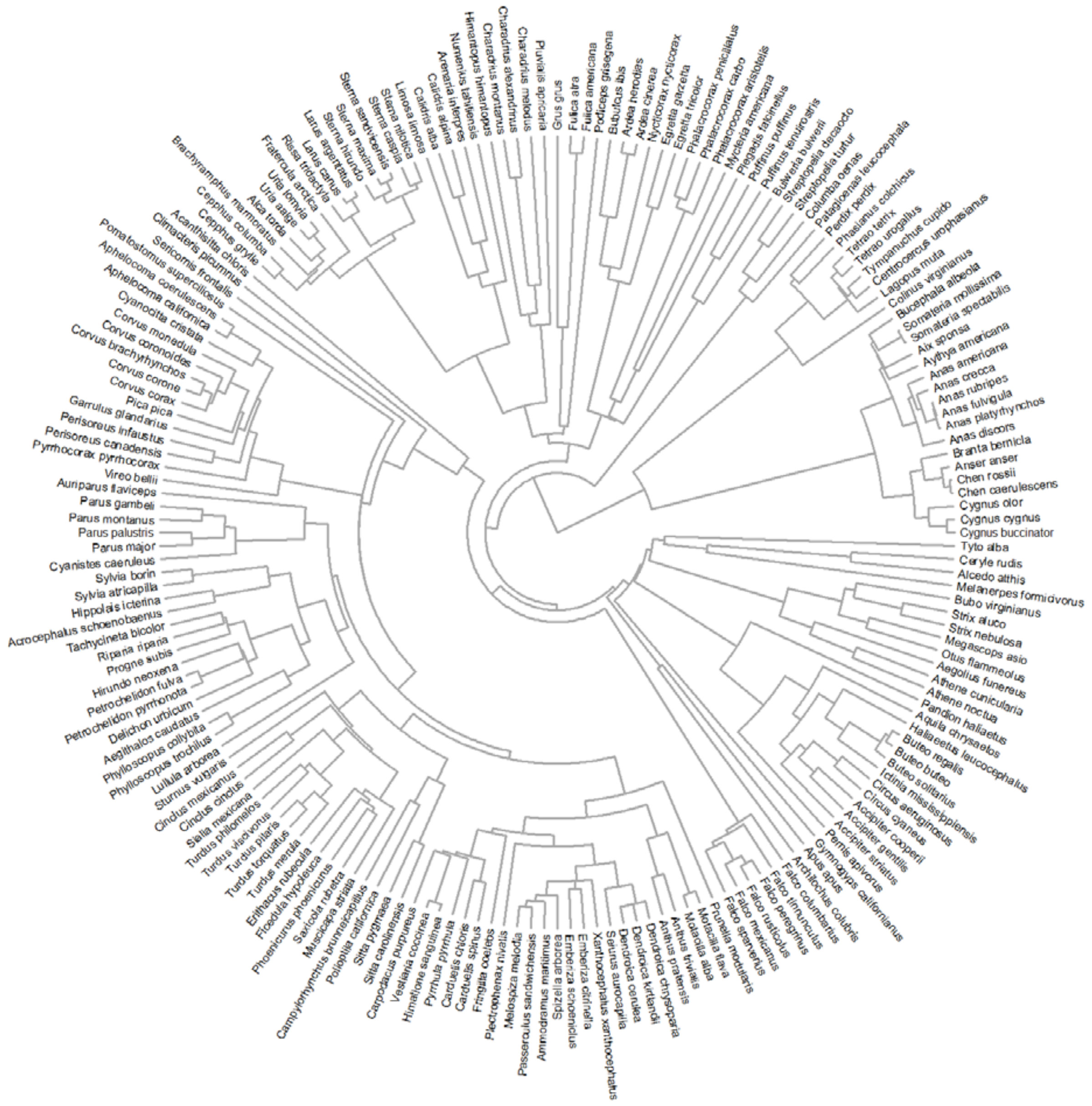


**Fig. S4.** Phylogenetic tree for the set of 188 species (based on the full tree from 1).

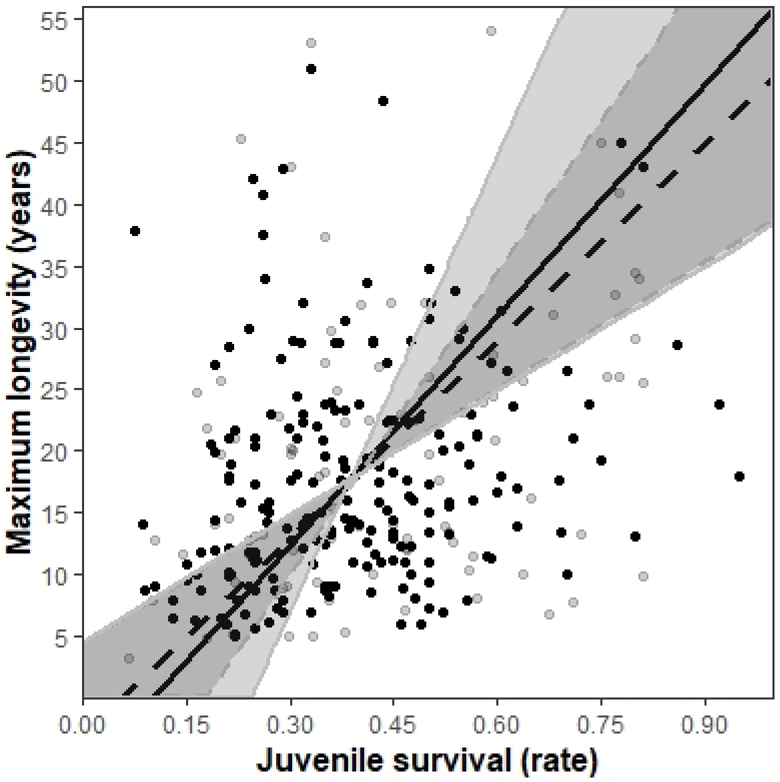


**Fig. S5.** Correlation between juvenile survival and maximum longevity on a) 293 species (all points, dashed regression line): RMA slope = 53.15, 95% CI (34.13, 81.71), $r_{Spearman} = 0.28$, S = 3003600, p <0.0001, and b) the 204 species included in the mixed model analyses (black point, black regression line): RMA slope = 62.18, 95% CI (33.52, 123.07), $r_{Spearman} = 0.27$, S = 1034800, p <0.0001. The shaded areas are the 95%CI of the RMA regression lines.

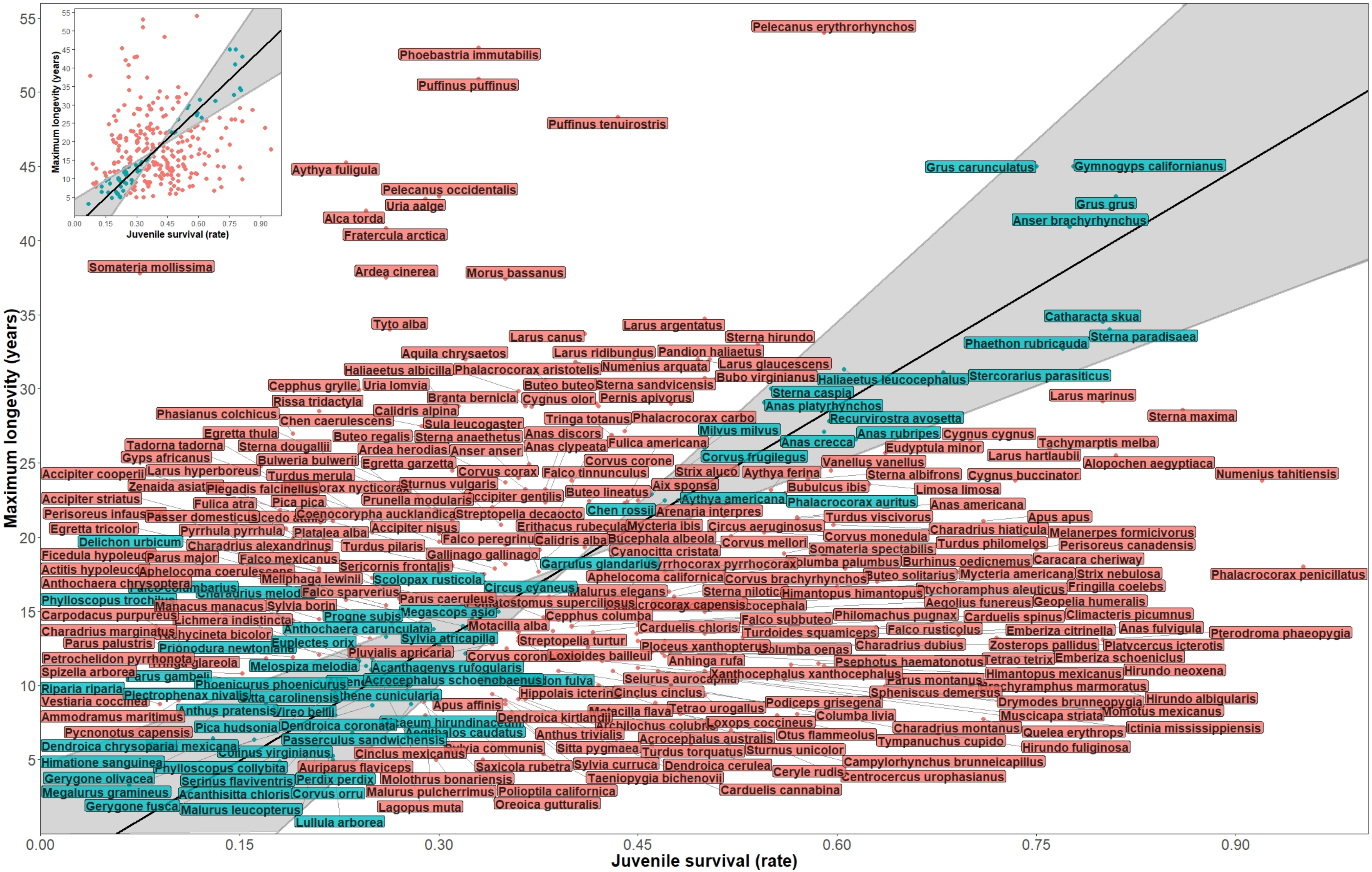


**Fig. S6.** Correlation between juvenile survival (first-year survival) and maximum longevity on 293 species with differentiation between the species inside (blue species; consistent with the ETA) and outside (pink species; contradicting the ETA) the RMA slope 95%CI (shaded area).

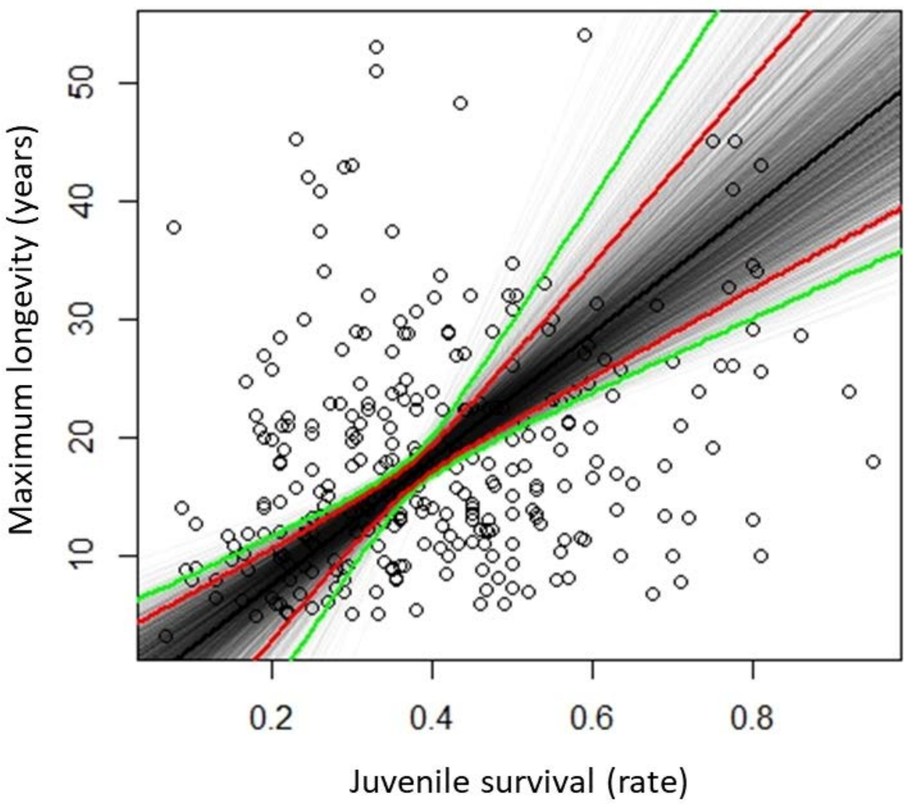


**Fig. S7.** Correlation between juvenile survival and maximum longevity on 293 species. RMA slope (in black), its 95% CI (in red: 78% species inside, 22% species outside) and its 99% CI (in green: 71% species inside, 29% outside). The grey lines depict patterns obtained for bootstrapping samples drawn from the data.

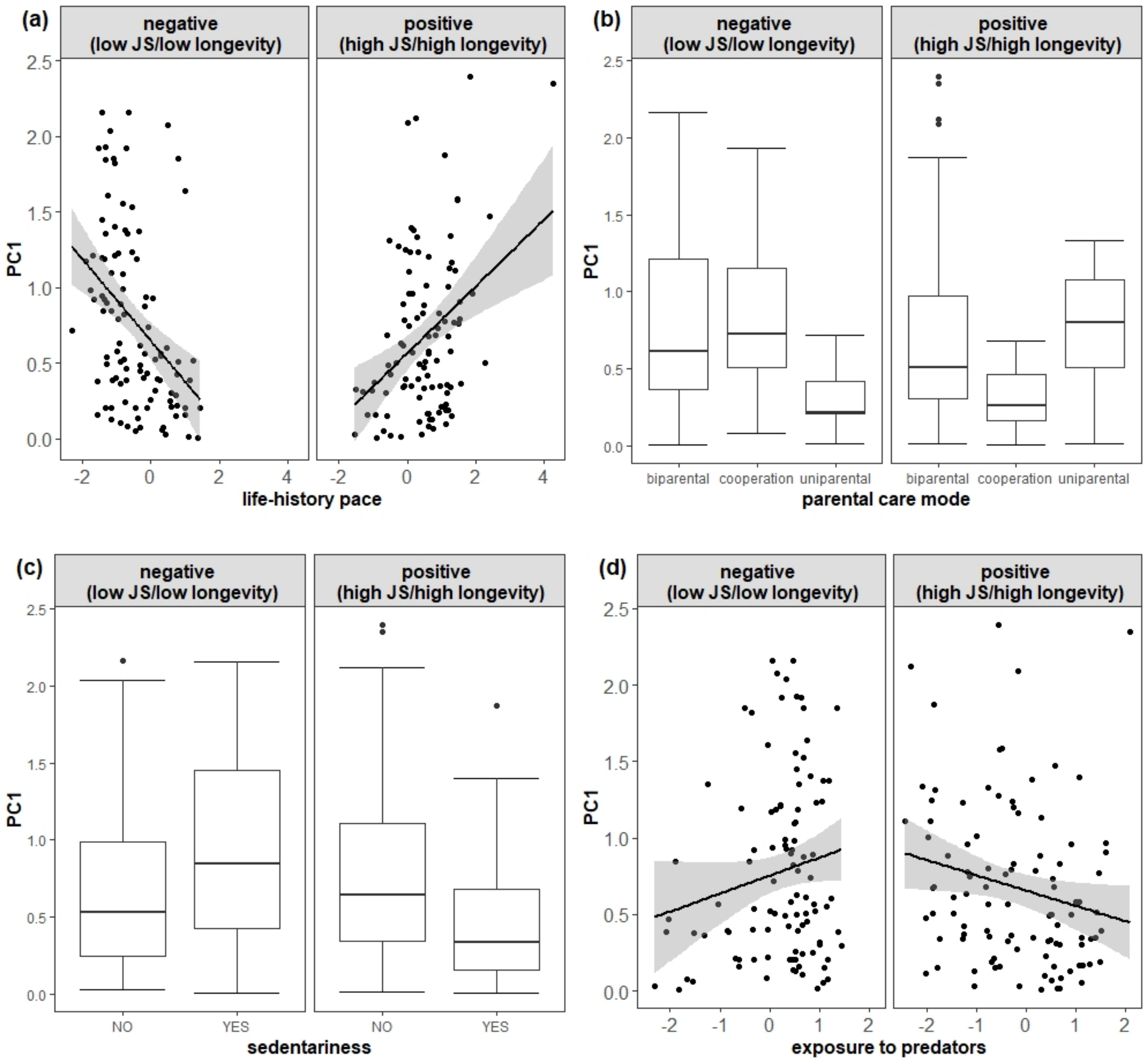


**Fig. S8.** Absolute PC1 in function of the sign of its raw values and of **(a)** life-history pace, **(b)** parental care mode, **(c)** sedentariness and **(d)** exposure to predators. JS = juvenile survival (first-year survival) (N=204).

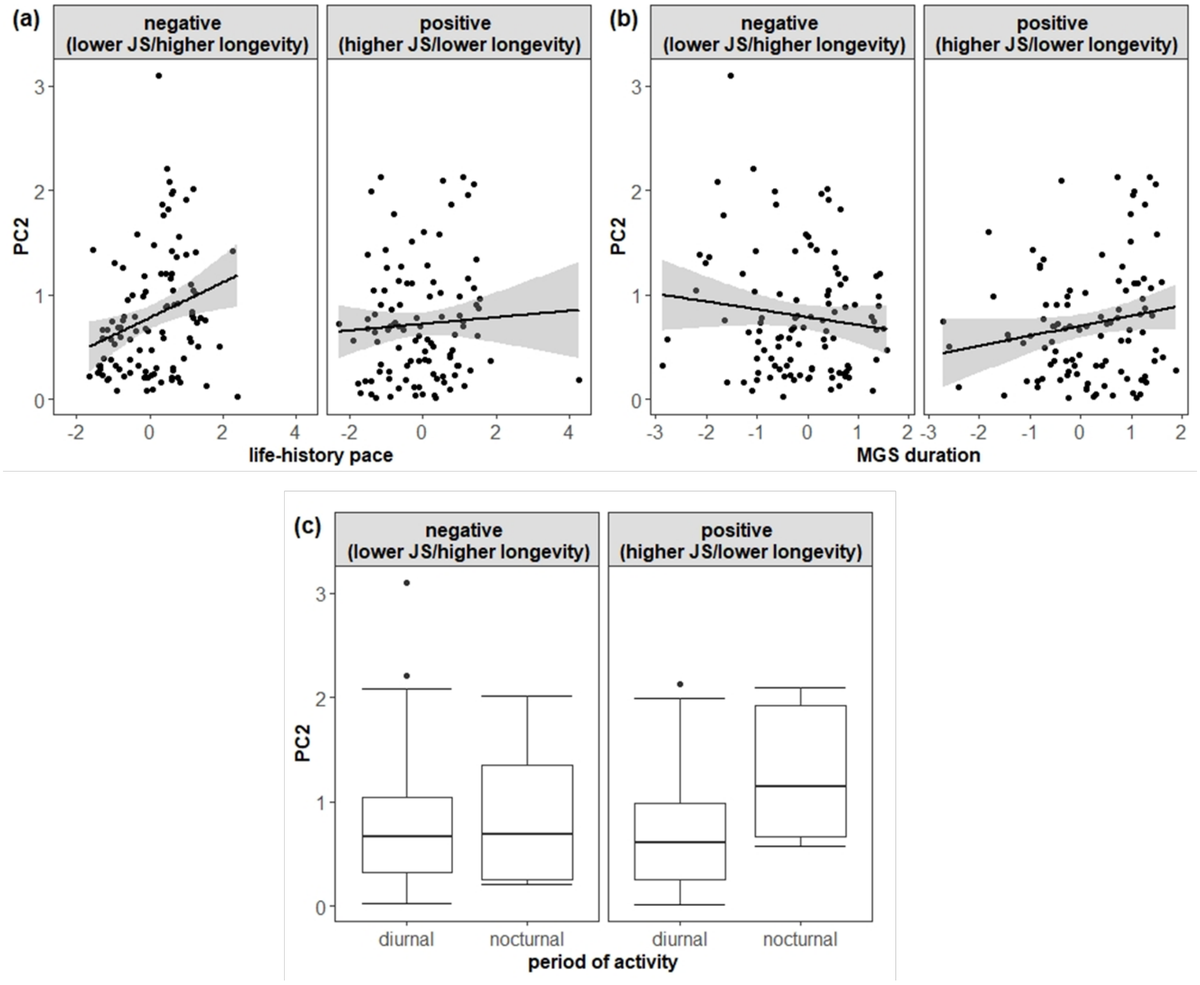


**Fig. S9.** Absolute PC2 in function of the sign of its raw values and of **(a)** life-history pace, **(b)** MGS duration and **(c)** period of activity. JS = Juvenile survival (first-year survival), MGS duration = mean duration of the growing season (N=204).

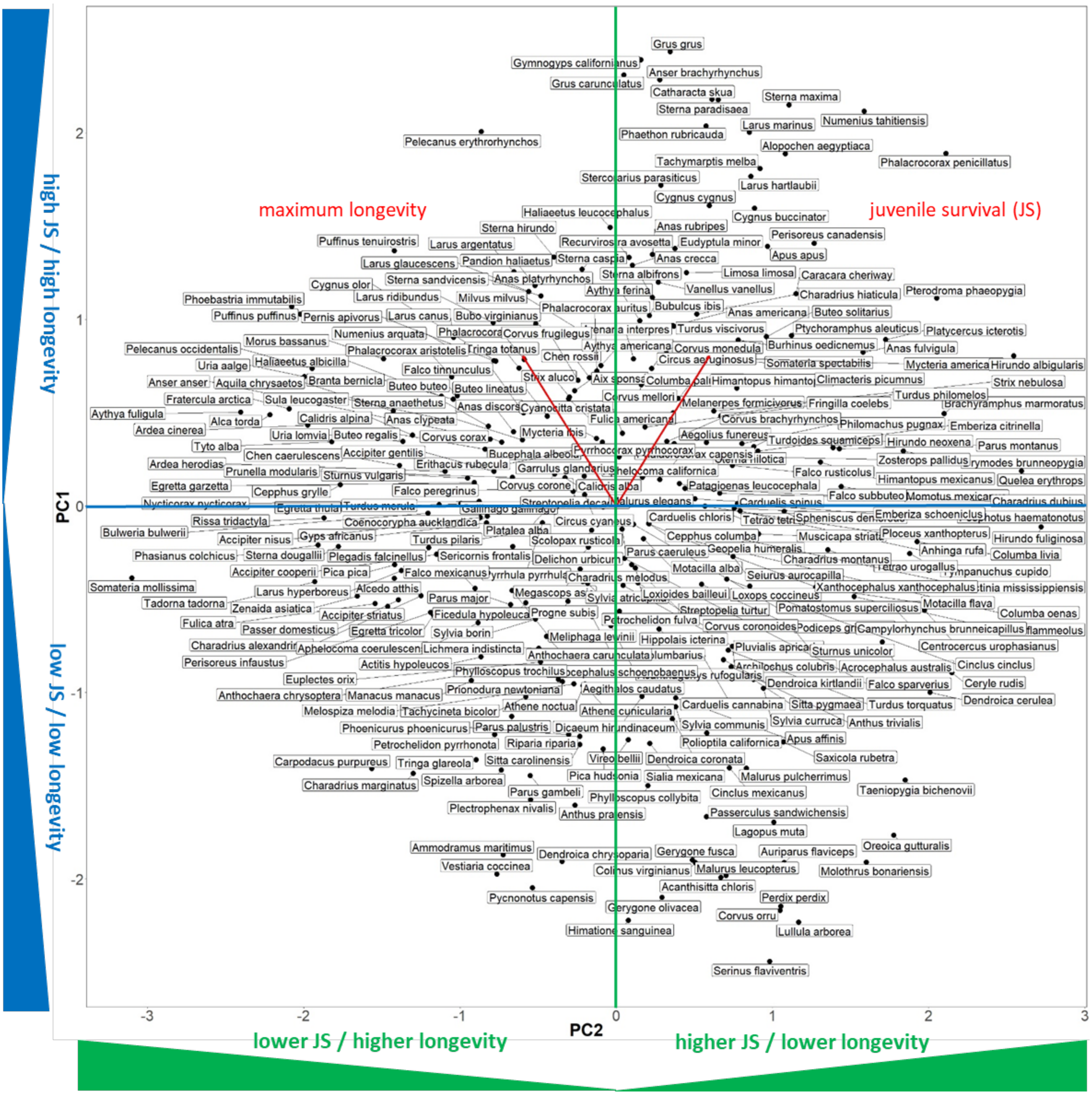


**Fig. S10.** Output of the PCA on juvenile survival (first-year survival, JS) and longevity with the indication of the species name.

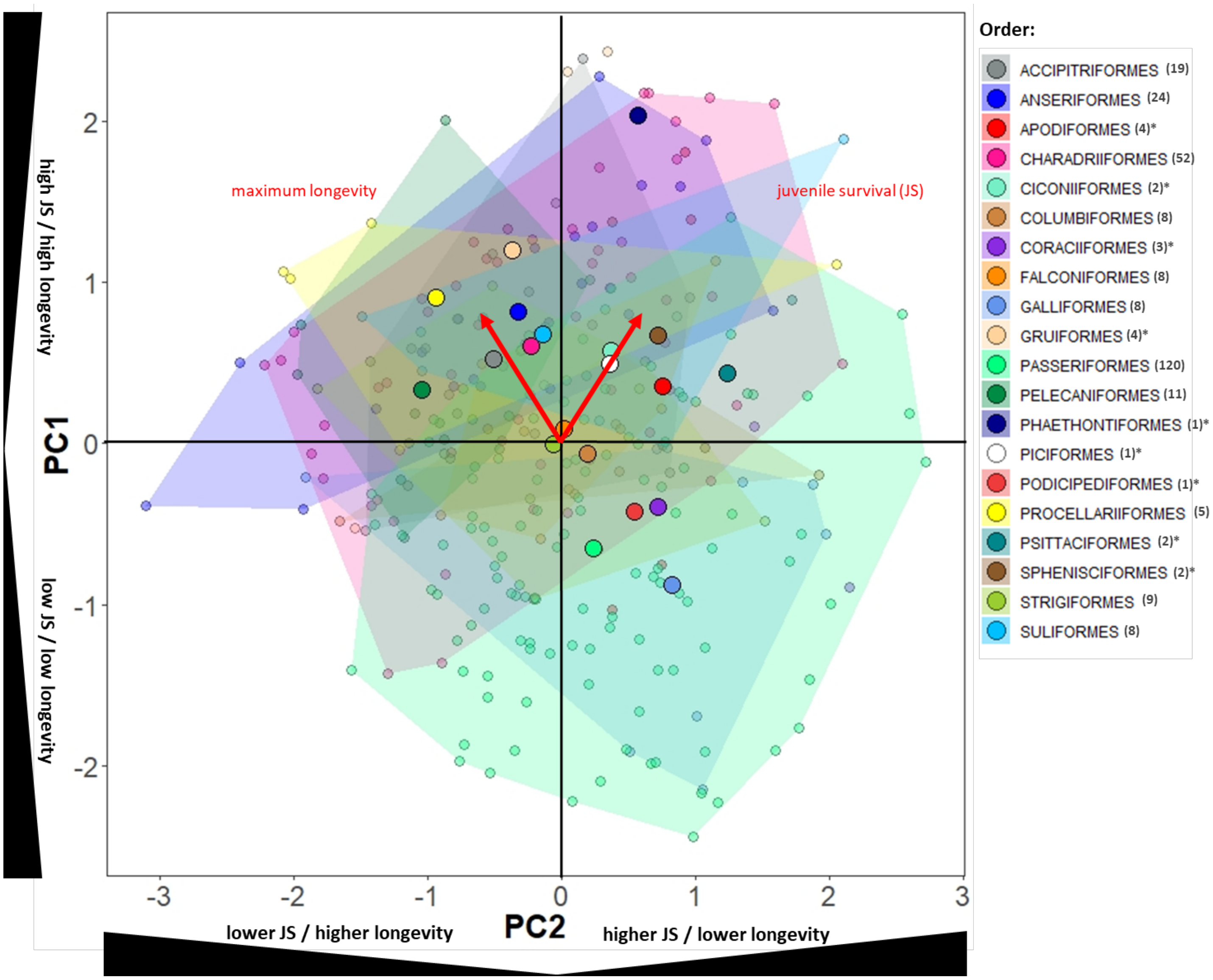


**Fig. S11.** Output of the PCA on juvenile survival (first year survival, JS) and longevity with the order identification (number of species per order given in the legend in parenthesis). (*) orders including less than 5 species for which the area is not drawn (for the purpose of illustration simplification).
